# HIV-1 expression is heterogeneous among clones of CD4+ T cells carrying authentic intact latent proviruses

**DOI:** 10.1101/2025.08.05.668730

**Authors:** Cintia Bittar, Ana Rafaela Teixeira, Thiago Y. Oliveira, Gabriela S. Silva Santos, Klara Lenart, Marcilio Jorge Fumagalli, Noemi L. Linden, Isabella A.T.M. Ferreira, Marina Caskey, R. Brad Jones, Mila Jankovic, Michel C. Nussenzweig

## Abstract

Antiretroviral therapy suppresses HIV-1 infection but is not curative because it fails to eliminate a reservoir of intact latent proviruses that reside primarily in CD4+ T cells. This compartment is composed of rare T cells that predominantly express memory and effector memory markers. The lack of precise understanding of the latent compartment has made it challenging to develop curative strategies for HIV-1 infection. Here we report on the properties of CD4+ T cells clones carrying intact latent proviruses, expanded *in vitro* from single cells obtained from the reservoir of people living with HIV-1. The latent proviruses in the clones were integrated into *ZNF* genes, non-genic satellite and centromeric regions, frequently associated with latency. Notably, the transcriptome of the cultured clones resembled their cells of origin. Despite their descent from single cells, only a fraction of the cells ranging from 0.4-14% expressed relatively low levels of HIV-1 that did not measurably alter host gene transcriptome. Latency reversing agents (LRAs) variably increased the number and amount of expression per cell, but the effects were modest and clone and LRA specific. The results suggest that pharmacologic and immunologic approaches to clear the reservoir should be optimized to accommodate intra- and inter-clonal diversity.

## Introduction

HIV-1 is an RNA virus whose life cycle requires reverse transcription and integration into the host genome. High level viral transcription is associated with cell death (Ruelas and Greene, 2013), and immunopathology. Anti-retroviral therapy (ART) prevents the spread of infection and suppresses viremia in people living with HIV-1 (PWH) but fails to cure the disease because intact proviruses persist in the genome of rare CD4+ T cells (Chun et al., 1998a; Finzi et al., 1997; Wong et al., 1997). This proviral reservoir is long-lived and responsible for rapid rebound viremia in individuals that interrupt therapy and it represents the primary barrier to HIV-1 cure (Cohn et al., 2020; Davey et al., 1999; Margolis and Archin, 2017; Siliciano and Siliciano, 2022).

Understanding the nature of the reservoir has been challenging because cells carrying intact latent proviruses are rare and there are no definitive markers that can be used to isolate them (Cohn et al., 2018; Collora et al., 2022; Crooks et al., 2015; Ho et al., 2013; Sun et al., 2023; Wei et al., 2023; Weymar et al., 2022; Wong et al., 2023; Wu et al., 2023). Moreover, defective proviruses are far more abundant than intact making it difficult to interpret simple proviral DNA measurements (Cho et al., 2022; Peluso et al., 2020). Quantitative estimates of the reservoir were initially made using *in vitro* outgrowth assays that depend on the ability of a given stimulus to induce proviral expression (Chun et al., 1998b; Finzi et al., 1997). These measurements were later shown to be underestimates because only a fraction of the latent cells were induced to produce virus after one round of stimulation (Ho et al., 2013; Hosmane et al., 2017). Nevertheless, outgrowth assays established that the reservoir decays rapidly in the first year after ART, but the rate of decay decreases to an estimated half-life of 4-7 years which explains why this therapy fails to eliminate the infection (Bachmann et al., 2019; Cho et al., 2022; Crooks et al., 2015; McMyn et al., 2023; Peluso et al., 2020; Siliciano et al., 2003).

Newer direct nucleic acid-based assays revealed that the proviral reservoir is primarily found in expanded clones of CD4+ T cells that wax and wane over time (Bui et al., 2017b; Cohn et al., 2015; Lorenzi et al., 2016; Lu et al., 2018; Simonetti et al., 2016). Sequencing revealed that the intact reservoir decays at an accelerated rate compared to the defective suggesting that there is either direct counterselection against intact proviruses by viral cytopathic effects, or immune-mediated elimination (Cho et al., 2022; Peluso et al., 2020; Reeves et al., 2023). Consistent with this idea, qualitative measurements of proviral mRNA expression showed that some latent proviruses are transcriptionally active (Dube et al., 2023; Einkauf et al., 2022; Procopio et al., 2015; Wiegand et al., 2017). Moreover, the relative rate of proviral transcription appears to be dependent on the position of the provirus in the genome (Battivelli et al., 2018; Collora and Ho, 2023; Jordan et al., 2001; Teixeira et al., 2024). Finally, prolonged therapy and elite control enrich for proviruses integrated into transcriptionally silent regions emphasizing the importance of proviral transcription on selection (Einkauf et al., 2019; Einkauf et al., 2022; Huang et al., 2021; Jiang et al., 2020).

One approach to accelerate reservoir elimination is to pharmacologically increase proviral transcription, by latency reversing agents (LRAs) (Archin et al., 2012; Armani-Tourret et al., 2024; Olesen et al., 2015; Rasmussen et al., 2014; Sogaard et al., 2015) Human trials using these agents can produce measurable increases in HIV-1 in circulation and changes in the population of CD4+ T cells carrying integrated proviruses. However, overall changes to reservoir size have been difficult to document possibly because of limited exposure or response heterogeneity within and between expanded clones of infected CD4+ T cells (Gunst et al., 2020; Tanaka et al., 2022).

Combinations of outgrowth and sequencing assays suggested that provirus transcription within a clone of CD4+ T cells can be heterogeneous (Einkauf et al., 2022; Hosmane et al., 2017). However, precise understanding of how transcription varies among cells within clones of CD4+ T cells bearing intact proviruses is limited. To further understand this phenomenon, we developed methods to identify and expand CD4+ T cells harboring intact latent proviruses in tissue culture (Teixeira et al., 2024; Weymar et al., 2022). Preliminary results indicated that the transcriptional profiles of the cultured cells resembled their *ex vivo* counterparts and that they expressed relatively low levels of HIV-1 (Teixeira et al., 2024; Weymar et al., 2022). Here we report on 6 cell lines derived from primary CD4+ T cells harboring intact proviruses, their heterogeneous levels of proviral expression, the effects of LRAs on the provirus and the impact of the provirus on the host cell transcriptome.

## Results

To obtain clones of CD4+ T cells harboring intact latent proviruses CD4+ CD45RA-T cells from PWH were enriched for the proviral clone of interest by cell sorting based on T cell receptor b-chain (TCRb) expression (Weymar et al., 2022). The TCR associated with clones obtained from participants 5104, 5125, 603, 9247 and B207 were known (Weymar et al., 2022). We determined the TCRb associated with clones from participants 5203 and P301 by flow cytometry using combinations of anti-Vb and Cb antibodies as previously described (Weymar et al., 2022) (Supplementary figure 1A). Sorted cells were plated at a density of 5 cells/well in the presence of feeder cells and stimulated with anti-CD3 and -CD28 plus IL-2 in the presence of anti-retroviral drugs to prevent spread of infection (Fig. 1A). After 3 weeks (resting) cultures were screened by ddPCR and confirmed by sequencing for the presence of intact proviruses. Positive cultures were further expanded for an additional 3 weeks for analysis. In total we obtained 6 expanded clones of CD4+ T cells harboring intact replication competent proviruses, and one carrying a defective provirus (P301^d^), each from a different donor (Figs. 1A and B, Supplementary Table 1). The fraction of HIV-infected cells in the final cultures used for analysis ranged from 16% to 100% (Supplementary figure 1C).

**Figure 1.**
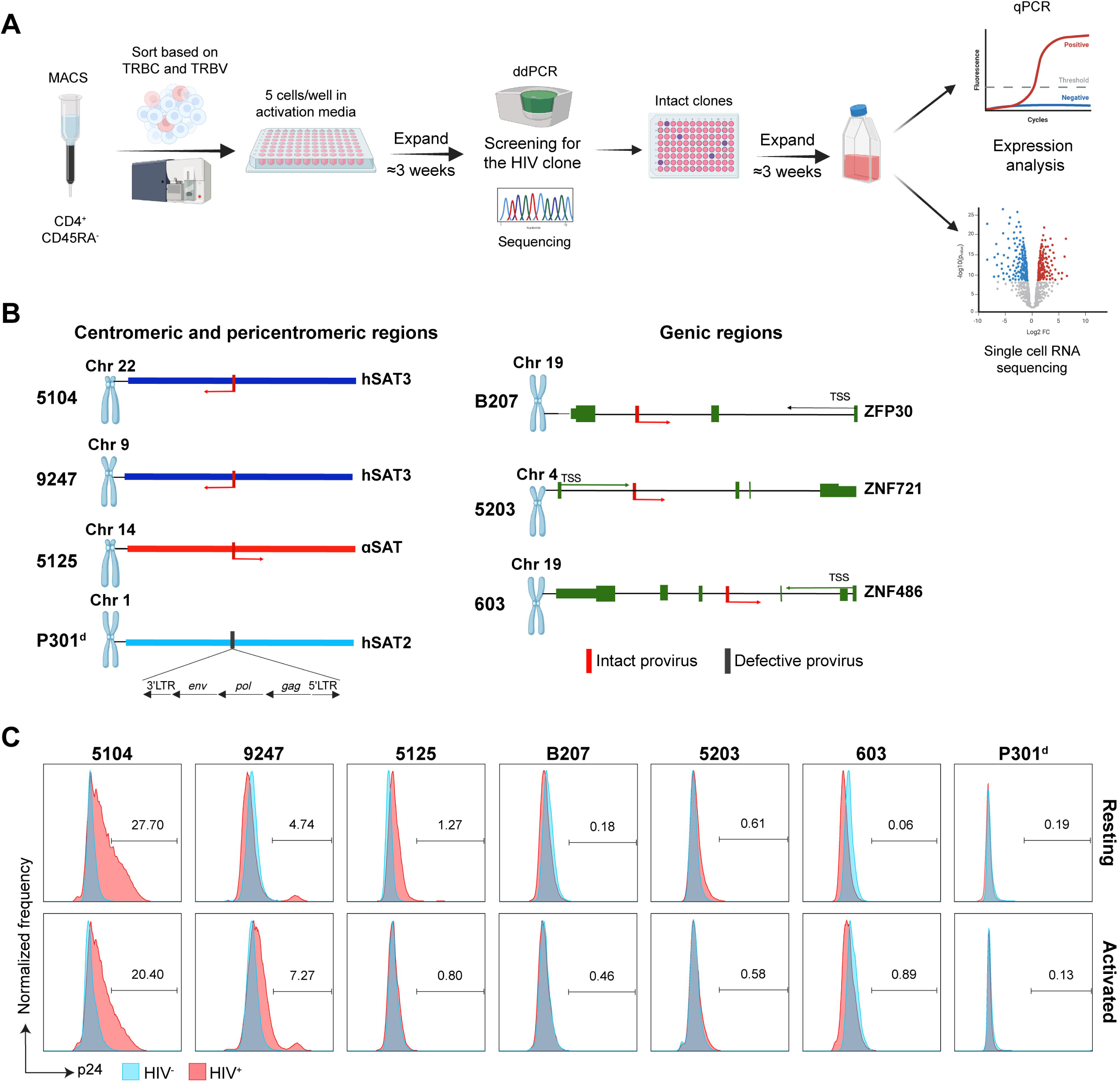
HIV-1+ CD4 T cells from PWH. **(A)** Graphic summary of the enrichment, screening, culture and analysis of the HIV-1+ CD4 T cell clones. **(B)** Provirus integration site for each clone. Arrows represent transcription orientation. Transcription from HIV-1 intact provirus is represented in red. The defective provirus is represented in black, with transcription arrows showing the location of the inversion. Transcription from host genes is represented in green. **(C)** Flow cytometry intracellular p24 detection in HIV-1+ CD4 T cell clones in resting and activated conditions Data from 5104 and 603 were previously presented in Teixeira et al. (2024).

The proviruses in 603, B207 and 5203 are integrated in the introns of genic regions of ZNF genes which are preferred sites of intact proviral integration (Einkauf et al., 2019; Einkauf et al., 2022; Huang et al., 2021; Jiang et al., 2020). In contrast, 5104 (Huang et al., 2021), 5125 and 9247 are in non-genic centromeric or pericentromeric satellite regions (Fig. 1B, Supplementary Table 1). P301^d^ is defective due to an inversion in its 5’ LTR, it is otherwise intact, and it too is integrated in a pericentromeric satellite region (Fig. 1B, Supplementary Table 1).

Primary cells corresponding to 4 of the clones (B207, 603, 9247, 5104) carry infectious proviruses (Cohn et al., 2018; Huang et al., 2021). To determine whether the 5125, 5203 and P301^d^ were able to produce infectious virus we used the supernatants of resting and anti-CD3 and -CD28 plus IL-2 stimulated (activated) cultures to infect HIV-1 negative donor CD4+ T cells. Six days after infection supernatants were assayed for HIV-1-Gag production by ELISA. HIV-1 Gag p24 was detected in the supernatants of 5125 and 5203 but not P301^d^, which is defective (Supplementary figure 1B). Thus, all but the defective clone can produce HIV-1. However, p24 expression as measured by flow cytometry in resting, or activated HIV-1+ CD4 cells was heterogenous (Fig. 1C).

To document HIV-1 transcription in the cultured cells we performed quantitative PCR (qPCR) for HIV-1 LTR, *gag* and *env.* We determined the number of mRNA copies using an HIV-1 standard and compared the amount of expression to cells productively infected with HIV-1_YU2_. The 6 clones that carry intact proviruses expressed different quantities of HIV-1, that were orders of magnitude lower than productively infected cells (Fig. 2A). LTR transcripts were barely detectable in 5203, 603 and B207 all of which are integrated into ZNF genes (Figs. 1B and 2A). Notably, higher levels of HIV-1 LTR expression were associated with intact proviruses integrated into pericentromeric satellite and centromeric regions (5104, 9247, 5125 Figs. 1B and 2A). *Gag* and *env* were proportional to LTR transcripts for all clones except P301^d^ that carries a defective provirus with an inverted 5’ LTR (Fig. 2A). Activation with anti-CD3 and -CD28 and IL-2 increased proviral expression in all clones carrying proviruses integrated into ZNF genes and 5125 which is integrated in a centromeric region (Fig. 2A, Supplementary Table 1). In conclusion mRNA expression data suggested that expanded clones of CD4+ T cells derived from a single provirally infected cell express low levels of HIV-1 (Einkauf et al., 2022) (Fig. 2A).

**Figure 2.**
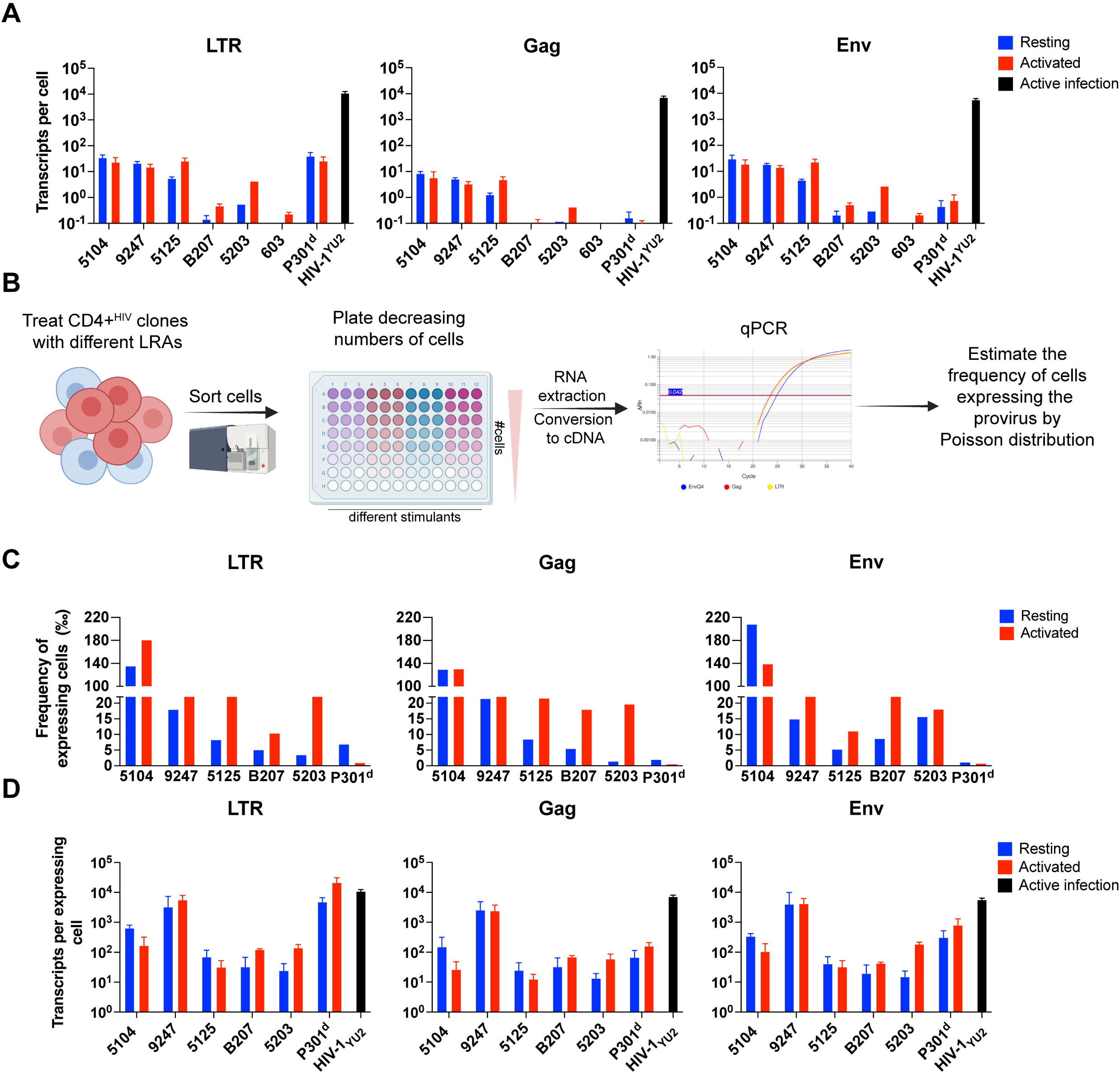
HIV-1 provirus expression in resting and activated conditions. **(A)** Histograms show average of HIV-1 transcripts per cell in resting (blue) and activated (red) conditions for the indicated clones and CD4^+^ T cells infected with HIV-1_YU2_ for comparison (black). Copies of LTR, *gag* and *env* transcripts were determined based on a standard curve. Average copy number per cell was calculated by dividing the estimated number of transcripts by the number of HIV+ cells in the assay. **(B)** Graphic representation of the limiting dilution assay. **(C)** Frequency of cells expressing HIV-1 LTR, Gag and Env transcripts in resting (blue) and activated (red) conditions. The number of expressing cells was estimated by Poisson distribution, based on limiting dilution data, calculated using the Most Probable Number (MPN) method. **(D)** Average number of transcripts per expressing cell in resting (blue) and activated (red) conditions, estimated based on the frequency of cells expressing HIV-1 transcripts as shown in “C”. Average transcripts per cell of an active infection by HIV-1_YU2_ is shown for comparison (black). All experiments were performed in triplicates. Frequency of cells expressing and copies of transcripts could not be calculated for 603 due to the low number of transcripts.

Flow cytometry and qPCR expression data suggest that expression of HIV-1 is heterogenous at the single cell level. To determine the fraction of cells within a clone that express proviral mRNA we performed limiting dilution experiments and measured HIV-1 transcripts by qPCR (Fig. 2B). Under resting conditions, the fraction of cells in a clone that expressed HIV-1 ranged from 4 to 140 per thousand (Fig. 2C). Activation with anti-CD3 and -CD28 increased the frequency of expressing cells in all cases tested except for P301^d^, that harbors a defective provirus (Fig. 2C). Notably, because the number of cells that expressed the provirus was low, the actual amount of HIV-1 mRNA per expressing cell could reach levels that were nearly comparable to that of a productive infection (Fig. 2D).

Proviral transcription can be modified by LRAs but little is known of their effects on single latent proviruses. To determine how proviral expression is modified by LRAs we exposed resting cells to these agents and measured HIV-1 LTR, *gag* and *env* expression in limiting dilution experiments by qPCR. Cultures were exposed to each of five different LRAs: Romidepsin, Panobinostat and Suberoylanilide hydroxamic acid (SAHA) which are histone deacetylase (HDAC) inhibitors; Prostratin, an NF-kB activator; and JQ1, an inhibitor of bromodomain and extra-terminal motif (BET) proteins which competes with the transcriptional inhibitor P-TEFb (Rodari et al., 2021). The effect of LRAs on HIV-1 gene expression was compared to resting and activated cells.

The response to the different LRAs varied and was clone specific with modest increases in the number of cells expressing HIV-1 (Fig. 3A). In no instance were all cells in a clone induced to express HIV-1. The greatest fraction of expressing cells, 27%, was achieved with Panobinostat in clone 5104 in which 14% of the cells expressed HIV-1 under resting conditions. However, Panobinostat was relatively ineffective in most of the other clones. Prostratin was unique in that it measurably increased the percentage of expressing cells in all clones tested but the increases were modest (Fig. 3A). Based on the frequency of cells expressing HIV-1, qPCR data was used to determine whether the LRAs increased the amount of expression per cell (Fig. 3B). For all clones, at least one of the LRAs increased the amount of proviral expression per cell over baseline (Fig. 3B). However, their effects varied among the clones with HDAC inhibitors, Romidepsin and Panobinostat being the most consistent activators among the LRAs. In conclusion, the relative proportion of cells expressing HIV-1 and the amount expressed per cell in response to LRAs is heterogeneous and clone and stimulus specific.

**Figure 3.**
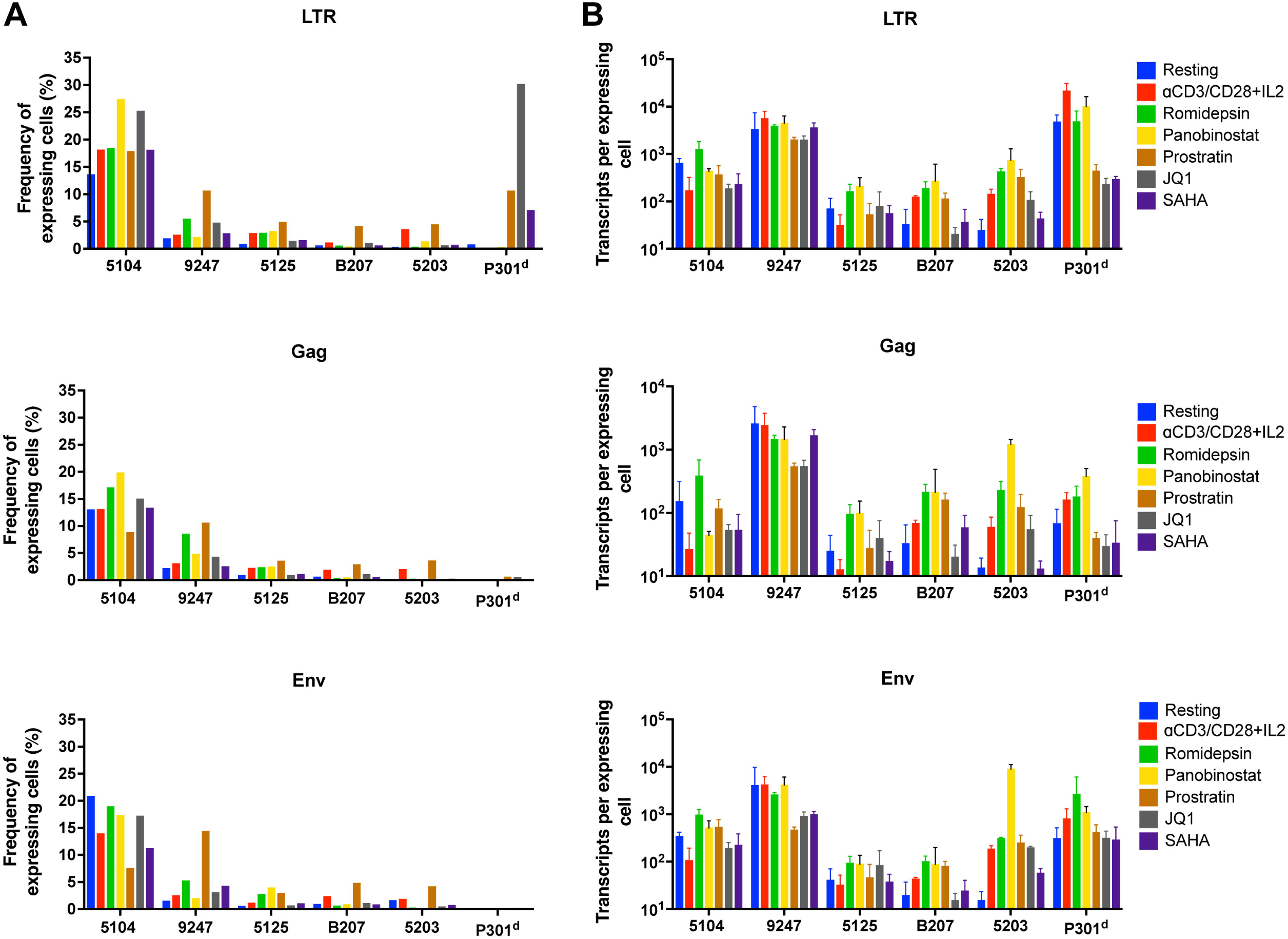
HIV-1 provirus expression after LRA treatment. **(A)** Frequency of cells expressing HIV-1 LTR, *gag* and *env* transcripts after treatment with LRAs estimated by poison distribution, based on limiting dilution data and calculated using the Most Probable Number (MPN) method. **(B)** Average copies of transcripts per expressing cell. The number of LTR, *gag* and *env* transcripts per reaction was determined based on a standard curve. Average copies of transcripts per expressing cell was estimated based on the frequency of cells expressing HIV-1 transcripts as shown in “A”. LRAs had no effect in HIV-1 provirus activation (data not shown) for clone 603. All experiments were performed in triplicates.

To examine the transcriptional profile of the cultured clones and how it compares to their primary *ex vivo* counterparts (Weymar et al., 2022) we performed single cell mRNA sequencing using the 10X Genomics platform on resting and activated cells. A total of 96,547 cells that carried an integrated provirus were analyzed (Supplementary figure 2 and Supplementary Table 2). TCR sequences were used to identify members of the CD4+ T cell clone of interest. Gene expression profiles were mapped onto primary cell data obtained from 5 of the 7 individuals from which clones were derived using a cut-off of 95% confidence (Fig. 4A). Although there was some heterogeneity in the primary cells and corresponding clones of CD4+ T cells, most of the clones mapped to similar or directly overlapping clusters (Fig. 4B). To determine which specific subset of T cells the clones were most like, we projected their gene expression profiles onto a reference multimodal single-cell data set of peripheral blood mononuclear cells (PBMCs) from HIV-1-negative individuals (Supplementary figure 3A). Consistent with the primary *ex vivo* data the clones were heterogeneous despite originating from single cells (Weymar et al., 2022). Most of the cultured clones showed a T-central memory phenotype, but others were composed of mixtures of CD4+ cytotoxic T lymphocytes (CTL) and a variety of different T-central and T-effector memory phenotypes (Supplementary figure 3B).

**Figure 4.**
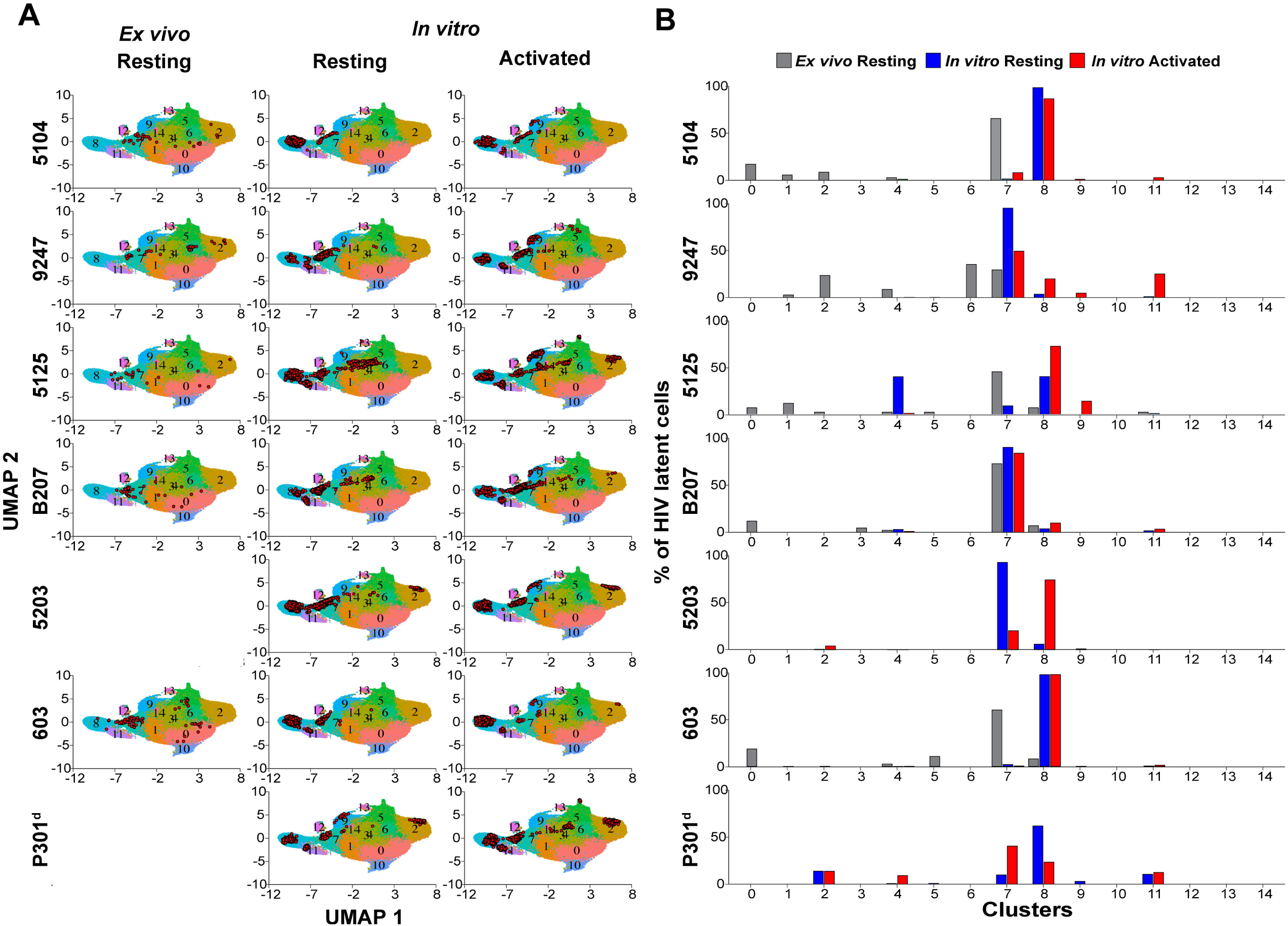
Uniform manifold approximation and projection (UMAP) of single-cell RNA sequencing data. **(A)** Uniform manifold projections (UMAPs) of gene expression profiles mapped on primary cell data based on 141,646 resting *ex vivo* cells from Weymar et al. (2022) and 96,547 *in vitro* cells (45,297 resting and 51,250 activated) from this study. The locations of cells from the HIV-1+ clone of interest are presented as red dot. **(B)** Bar graph representing the fraction of cells that fall into each of the 15 clusters from the UMAP. *Ex vivo* resting cells are presented in gray, *in vitro* resting cells in blue and *in vitro* activated cells in red. All experiments were performed in duplicate.

To determine whether there are unique transcriptional features shared by CD4+ T cells clones that harbor latent proviruses we compared their transcriptomes to non-infected cells within the same cultures (Supplementary Table 3). Each of the clones differed significantly from the uninfected co-cultured controls (Supplementary Table 3). To determine whether the differentially expressed genes are shared by different clones we compared the 6 intact and one defective clone. We found 41 and 20 differentially upregulated genes that were shared among at least 4 clones in resting and activated conditions respectively (Fig. 5). Among these 17/41 and 13/20 differentially upregulated genes from resting and activated cells respectively were previously identified as enriched among CD4+ T cells harboring intact latent proviruses (Figs. 5A and 5B). The differentially expressed genes included *HLADR* genes, *CCL5* and genes expressed by cytotoxic T cells, consistent with previous reports (Cohn et al., 2018; Collora et al., 2022; Weymar et al., 2022). In conclusion the cultured clones resemble their primary *ex vivo* counterparts.

**Figure 5.**
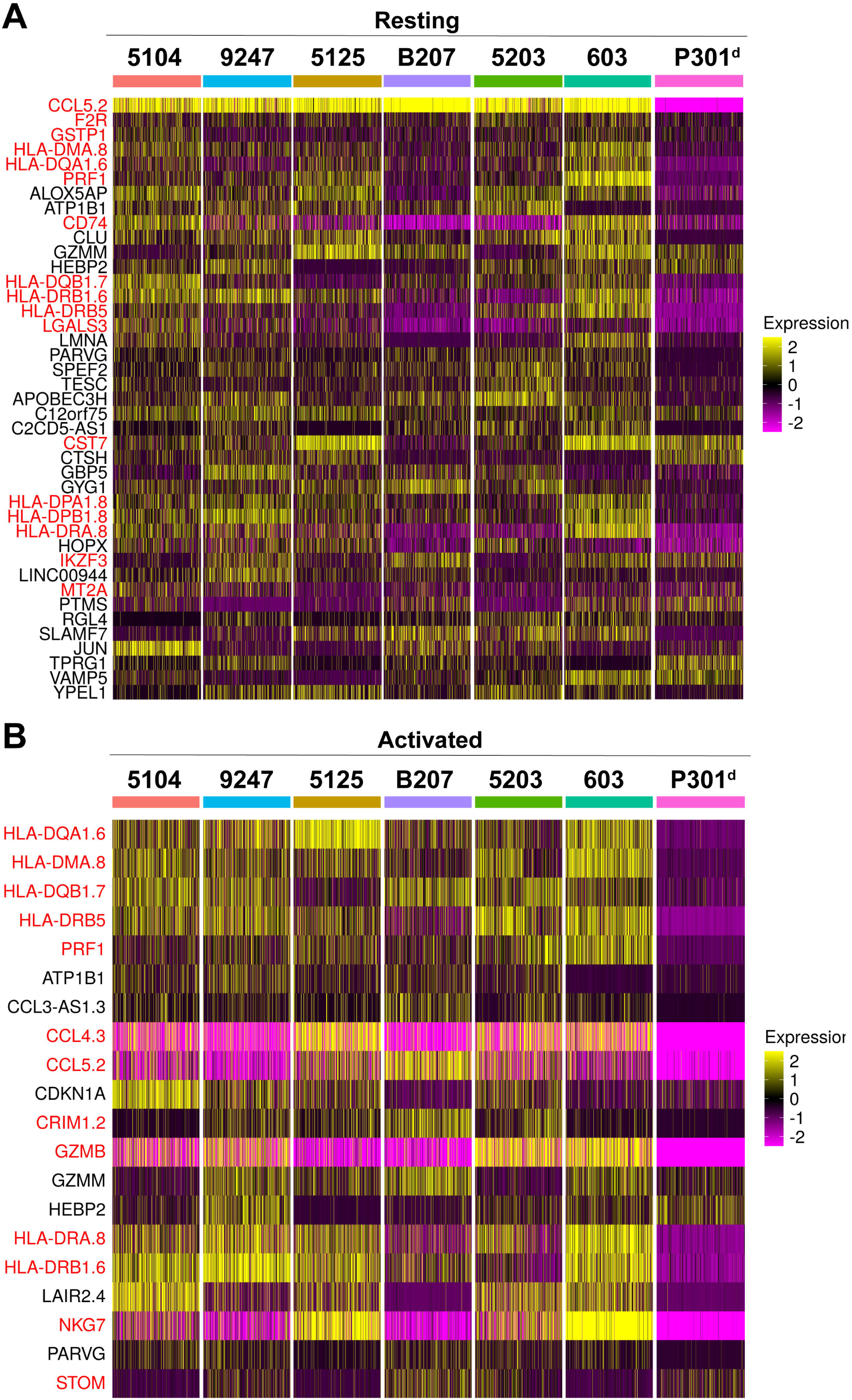
Overlapping DEGs between HIV-positive and HIV-negative CD4+ T cells. Single-cell heatmaps showing mRNA expression of overlapping differentially expressed genes that are upregulated in HIV-positive cells compared to HIV-negative cells (rows), detected in a minimum of 10% of cells from at least four clones under resting **(A)** or activated **(B)** conditions. A subset of cells from each clone (columns) was selected for visualization.

Heterogeneous HIV-1 expression was observed by flow cytometry and qPCR and confirmed by single cell RNA sequencing which showed that despite their single cell origin only a fraction of cells in a clone express HIV-1 (Supplementary table 4). To determine whether HIV-1 expression by members of a clone alter cellular transcription we compared the transcriptome of individual cells within a clone that express the LTR and those that do not (Fig. 6A and Supplementary figure 4). Notably, HIV-1 transcripts were the only differentially expressed mRNAs shared by all clones (Fig. 6B). We conclude that the HIV-1 expression in clones of CD4+ T cells harboring latent proviruses do not induce a common cellular transcriptional response.

**Figure 6.**
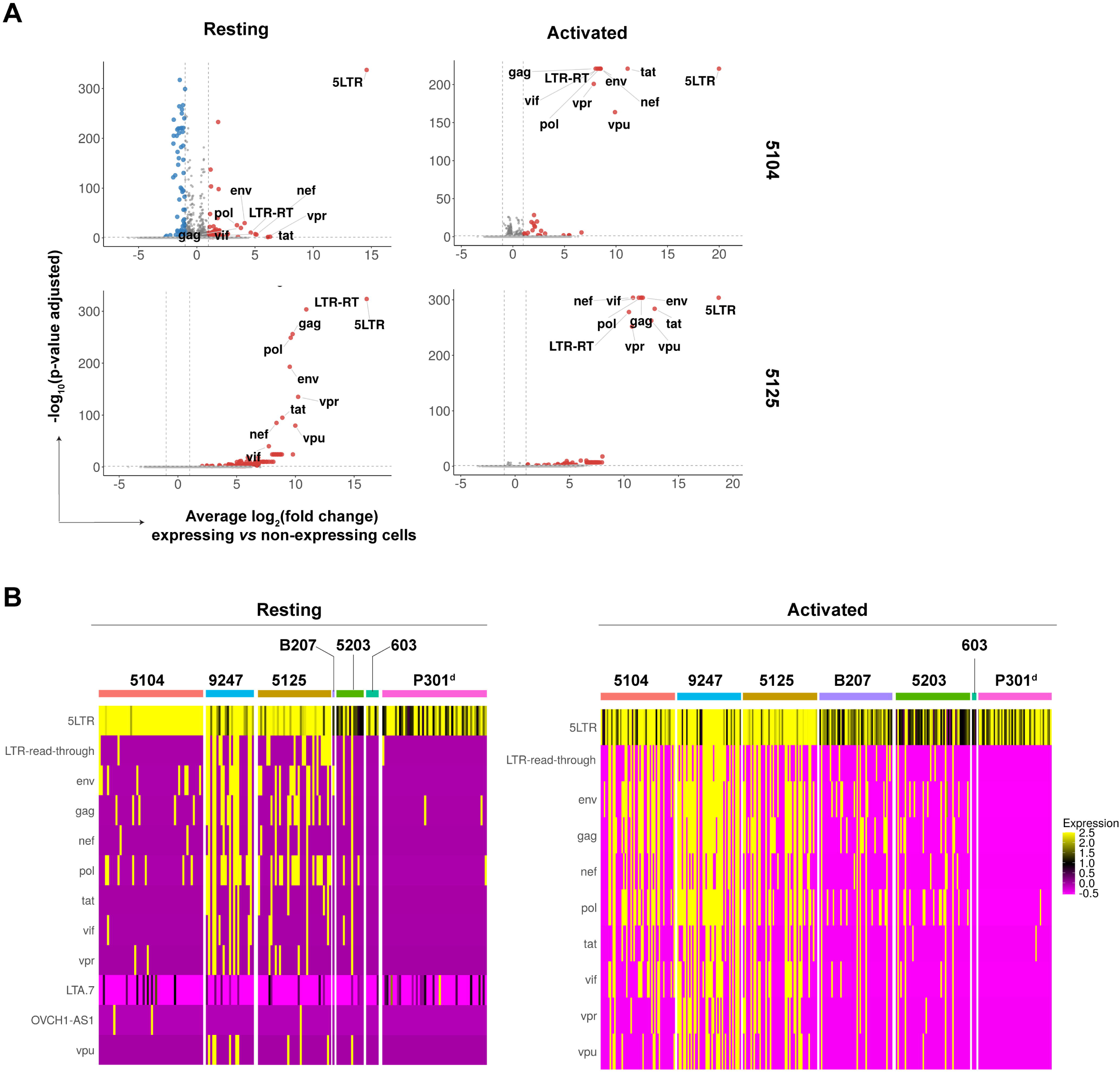
Differential expression between HIV-infected cells positive and negative for viral transcripts. **(A)** Volcano plots showing differentially expressed genes between HIV-expressing and non-expressing cells from clones 5104 and 5125 under resting or activated conditions. **(B)** Heatmap showing expression of overlapping differentially expressed genes between HIV-expressing and non-expressing cells under resting or activated conditions. A subset of cells from each clone (columns) was selected for visualization.

## Discussion

Experiments in animal models, observational studies in humans that spontaneously control infection, and small interventional clinical studies indicate that the immune system can durably control infection. In HIV-1 controllers and PWH on long term therapy the proviruses that remain in the reservoir tend to be integrated into transcriptionally silent parts of the genome suggesting that strategies that increase proviral transcription and enhance immune responses would help eliminate the reservoir (Einkauf et al., 2019; Einkauf et al., 2022; Huang et al., 2021; Jiang et al., 2020; Lian et al., 2023). However, the limited clinical studies performed to date show little or no measurable effect of LRAs on the HIV-1 reservoir despite documented effects in proviral transcription (Debrabander et al., 2023). In this study we examined intact and defective proviral transcription by clones of CD4+ T cells derived from single cells from 7 different ART suppressed individuals. Our observation that baseline and LRA stimulated levels of proviral transcription vary within cells in a latent clone and between clones suggests that combinations of these agents would need to be used to optimize clinical efficacy.

The cells that we selected for expansion *in vitro*, were derived from PWH suppressed on ART. As might be expected, the proviral integration sites were biased to ZNF genes and non-genic centromeric and satellite regions both of which are poorly transcribed (Cano-Gamez et al., 2020; Einkauf et al., 2022; Jordan et al., 2003; Lewinski et al., 2005). The somewhat higher levels of HIV-1 transcription among proviruses integrated in centromeric regions than in ZNF genes may be due to constitutive low level RNAPol-II dependent transcription of those parts of the genome (Perea-Resa and Blower, 2018; Zhu et al., 2023).

Although not entirely identical, the cells that we expanded *in vitro* are closely related to their *ex vivo* counterparts and typically show a T cell memory or effector memory phenotype (Weymar et al., 2022). Moreover, the genes that were differentially expressed between provirus-containing cells and their uninfected co-cultured counterparts were many of the same genes differentially expressed by latently infected primary cells (Cohn et al., 2018; Collora et al., 2022; Horsburgh et al., 2020; Sun et al., 2023; Weymar et al., 2022). Our experiments demonstrate that the difference between infected and uninfected cells cannot be attributed to cellular response to proviral expression because the only reproducible difference between HIV-1 expressing resting or activated cells within a latent clone is HIV-1. Thus, the differences are likely due to yet to be determined factors that impact T cell infectivity and/or fate decisions at the time of or shortly after exposure to the virus.

Over 50% of intact latent proviruses found in PWH reside within expanded clones of CD4+ T cells suggesting that their proliferative expansion occurs in the absence of dominant cytopathic effects (Bui et al., 2017b; Cohn et al., 2015; Hosmane et al., 2017; Lorenzi et al., 2016; Simonetti et al., 2016). Like uninfected cells, CD4+ T cells bearing intact latent proviruses expand in response to antigen including CMV, EBV and HIV-1, antigens associated with chronic infections (Demoustier et al., 2002; Douek et al., 2002; Niessl et al., 2020; Simonetti et al., 2021). Theses clones survive long term and proliferate *in vivo*, and their component cells can produce infectious virus particles when stimulated *in vitro*. Anti-CD3 and CD28 plus IL-2 stimulation mimics strong T cell activation signals. Notably stimulation of mixtures of primary cells *in vitro* only induces virus production by some latent cells (Bui et al., 2017a; Hosmane et al., 2017). One potential explanation for these results is that transcription of the provirus within expanded clones obtained from PWH is heterogeneous with some cells producing virus while others remain silent after stimulation. Our experiments support this idea and demonstrate that even after extensive rounds of division *in vitro* under strong stimulating conditions intact latent proviral transcription is heterogeneous within a clone of CD4+ T cells derived from a single primary cell.

The heterogeneity of HIV-1 expression in clones reflects a variety of stochastic mechanisms that normally regulate cellular transcription. These include binding kinetics of transcription factors, enhancer interactions with gene promoters and chromatin architecture (Brouwer and Lenstra, 2019). Thus, intra-clone variability in HIV expression can be attributed to stochastic cellular transcription dynamics in the region of proviral integration combined with complex regulation of the HIV-1 promoter (Damour et al., 2023; Tantale et al., 2021; Weinberger et al., 2005). Moreover, the observation that HIV-1 transcription in expanded clones of CD4+ T cells carrying authentic latent proviruses is stochastic is entirely in keeping with experiments in cell lines carrying randomly integrated indicator proviruses (Jordan et al., 2003; Lewinski et al., 2005).

Expanded clones of CD4+ T cells dominate the circulating intact proviral reservoir, but the bigger the clone the less likely it is that the virus contributes to rebound viremia when ART is interrupted (Lorenzi et al., 2016). There are several possible explanations for this phenomenon including immune selection against clones that produce high levels of virus, and suppression by autologous antibodies (Bertagnolli et al., 2020; Esmaeilzadeh et al., 2023; Lorenzi et al., 2016). An additional possibility suggested by our quantitative data is that although clones can be very large, the number of cells in a clone that produce virus and the amount of virus they produce at any time is small and therefore unlikely to contribute to rebound.

Targeting the HIV-1 reservoir for elimination by pharmacological methods or immune intervention has been challenging. Within clone and between clone heterogeneity in proviral expression at baseline and in response to LRAs represent important temporal hurdles to HIV-1 remission or cure. Our experiments help understand the nature of some of these impediments and suggest how pharmacologic and immunologic approaches to cure or remission should be optimized including longer lasting and combination interventions to maximally impact diverse clones and clonal members.

## Methods

### Study participants and samples

Samples used in this study originate from people living with HIV (PWH) who enrolled in clinical trials or a sample collection protocol at the Rockefeller University Hospital, New York, NY (NCT03526848: 5104, 5125, 5203, NCT02825797: 9247, NCT02588586: 603, protocol MCA-0966: B207) and at the University of Pennsylvania, Philadelphia, PA (NCT05245292: P301) (Cohn et al., 2018; Gaebler et al., 2022; Mendoza et al., 2018). Informed consent was obtained and samples were investigated under the Rockefeller University IRB-approved protocols TSC-0910 and MCA-0966. Total peripheral blood mononuclear cells (PBMCs) were isolated from leukapheresis by Ficoll separation and frozen in aliquots. All samples used in this study were collected at baseline time points of the clinical trials, except for sample 9247, for which the 12-week time point was used.

### Cell sorting and culture

CD4+CD45RA^-^ T cells were isolated from PBMCs by negative selection using magnetic separation (Miltenyi cat. 130-096-533; cat. 130-045-901). Fc receptor was blocked by incubating cells with Fc-blocking reagent (Miltenyi, cat. 130-059-901). CD4+CD45RA^-^ T cells were stained for live/dead detection at 1:1000 with Fixable Viability Dye eFluor 780 (Invitrogen, cat. 65-0865-14) and Brilliant Violet 605 anti-human TCR Cb1 (BD, cat. 747979), PerCP/Cy5.5 anti-human CD4 (BioLegend, cat. 317428), Pacific Blue anti-human CD3 (BioLegend, cat.300431) 1:100. Cells were also stained, at 1:100, with antibodies binding to TCRβ variable chains specific to the clone: FITC anti-human TCR Vβ17 (603) (Beckman Coulter, cat. IM1234); FITC anti-human TCR Vb7.2 (9247) (Beckman Coulter, cat. B06666); FITC anti-human TCR Vβ22 (5125) (Beckman Coulter, cat. IM1484) (Weymar et al., 2022). Beta Mark TCR Vbeta Repertoire Kit (Beckman Coulter, cat. IM3497) is a panel of 24 different anti-TRBV antibodies that are divided into 8 vials, with a mix of three 3 antibodies each (A to H), that are conjugated with PE, or FITC, or PE-FITC. Since there is no available commercial antibody for the TCRβ variable chains for clones 5104 (TRBV5-4) and B207 (TRBV7-8), we enriched by labeling with all antibodies from Beta Mark TCR Vbeta Repertoire Kit and sorted for the negative population. Individual vials from this panel were also used to stain cells from 5203 and P301, mix D (FITC+ TRBV 12-3+) and mix C (FITC-PE+ TRBV5-1^+^), respectively. In summary, latent cells’ enrichment was done by sorting the following CD4+CD3+ cell populations: B207 and 5104 - TRBC1+TRBV; 603 - TRBC1+TRBV19+; 5125 - TRBC1+TRBV2+; 9247 - TRBC1-TRBV4-3+; 5125 - TRBV12-3+; and P301 - TRBC1-TRBV5-1+.

Five cells per well were sorted by FACSymphony S6 using FACSDiva software (BD Biosciences version 9.5.1) in 96 well U-bottom culture plates. The plates contained 200µL/well of activation media composed of R10 (RPMI-1640 supplemented with 10% heat inactivated FCS, 10 mM HEPES, 100 U/ml penicillin–streptomycin and 2mM L-glutamine - Gibco) supplemented with 50 U/ml of recombinant human IL-2 (Roche cat. 10799068001), anti-human CD3 and CD28 antibodies (Biolegend cat. 317325; 302933), both at 100 ng/mL, and feeder cells (5000 rads irradiated PBMCs, depleted of NK and CD8+ T cells) at a concentration of 1 x 10^6^ cells/mL. To prevent the infection of new cells four antiretroviral (ARV) drugs were added to the media: Tenofovir (1 µM), Emtricitabine (1 µM), Nevirapine (1 µM), and Enfuvirtide (10 µM). The number of sorted plates for each clone was determined based on the predicted frequency in the enriched population (Weymar et al., 2022). Cells were cultured in a humidified incubator at 37°C with 5% CO^2^ for 3 to 4 weeks, with media exchange twice a week (maintenance media - R10 + IL-2 + ARVs). Cultures containing latent cells were further expanded in activation media.

### HIV provirus screening and enrichment

The presence of HIV-1 positive cells in the cultures was determined by digital droplet PCR (ddPCR) as previously described (Bruner et al., 2019). For DNA extraction, 2μL of the cultures were collected and transferred to a 96 wells PCR plate containing 8μL of RLT buffer (Qiagen cat 79216). Agencourt RNA Clean XP magnetic beads (Beckman Coulter cat. A63987) was used for DNA purification, according to manufacturer’s instructions. ddPCR reaction droplets were generated using the Automated Droplet Generator (Bio-Rad), reactions were read by QX 200 Droplet Reader (Bio-Rad) and data was analyzed using QX Manager Standard Edition software (version 2.0.0.665 Bio-Rad).

Env and near full length proviral PCR (NFL-PCR), followed by sanger sequencing (GeneWiz, Azenta Life Sciences)) and next-generation-sequencing (NGS) (MiSeq Nano V2 300 cycles, Illumina) respectively, were used to identify and characterize the provirus integrated in the CD4+ T cell clones, as described previously (Weymar et al., 2022).

### Integration site

Integration sites from the clones 603, 5104, 5125, B207, 9247 and P301 were determined or confirmed using the integration site loop amplification (ISLA) method, as previously described (Wagner et al., 2014). Genomic DNA was pre-amplified using multiple displacement amplification (Qiagen REPLI-g single cell advanced kit) prior to the ISLA reaction. Integration site from clone 5203 was confirmed by amplification of the integration site using HIVLTR1 primer and ZNF721Int primer based on available information (Supplementary table 5) (Huang et al., 2021).

### CD4+ T cell activation and latency reversal agents (LRA)

HIV-1 expression by CD4+ T cell clones was analyzed in resting (R10) and 24 hours after activation (50U/mL IL-2 + 100 ng/mL anti-CD3 and CD28 antibodies) and treatment with: romidepsin (5 nM, Active Motif cat. 14083), panobinostat (50 nM, Sigma-Aldrich cat. SML3060-10MG), prostratin (2.5 μM, Sigma-Aldrich cat. P0077-1MG), JQ1 (1 μM, Sigma-Aldrich cat. SML1524-5MG), SAHA (Sigma-Aldrich 0.5 μM, cat. SML0061-5MG). All assays were performed in the presence of ARVs.

### Flow cytometry

The expression of HIV-1 Gag p24 protein was analyzed, in resting and activated cells, by flow cytometry. Cells were fixed and permeabilized with BD Cytofix/Cytoperm Fixation/Permeabilization Kit (BD Biosciences cat. 554714) and stained with Gag p24-FITC-conjugated antibody (HIV-1 core antigen-FITC, clone KC57, Beckman Coulter cat. 6604665) diluted 1:100. Readings were performed in FACS Symphony A5 flow cytometer (BD Biosciences) running FACS Diva software version 8.5. Data were analyzed using using FlowJo software (version 10.10.0, BD Biosciences)

### HIV-1 Expression

To evaluate HIV-1 expression in bulk cultures cells were plated at a density of 2 x 10^5^ per well in a 96 wells plate. 24h after stimulation cells were counted, spun at 500g for 5 minutes to remove the supernatant and lysed in 350μL of RLT buffer (Qiagen cat 79216). Lysed samples were stored at -80°C. RNA was extracted using the RNeasy Plus Micro kit (Qiagen cat. 74034) according to manufacturer’s instruction. cDNA was synthesized using SuperScript III Reverse Transcriptase (Invitrogen cat. 18080-093) according to manufacturer’s instructions and, 2.5 ng/μL random primers (Invitrogen cat. 48190011) along with primers specific (0.1μM each) for long LTR, *gag*, *pol*, *tat-rev*, *nef*, *poly-A* viral transcripts (Einkauf et al., 2022; Yukl et al., 2018) (Supplementary table 5).

To estimate the frequency of cells expressing HIV-1 and the level of expression per cell, we performed limiting dilution assays. Defined numbers of cells were sorted into 96 well PCR plates, containing 10μL to 20μL of RLT Buffer (Qiagen cat. 79216) per well in triplicate. The number of cells sorted into individual wells was determined based on prior PCR estimates. RNA was purified with Agencourt RNACleanXP magnetic beads (Beckman Coulter cat. A63987), as above. To eliminate DNA, samples were eluted in 10μL DNAseI mix (Invitrogen cat. 18068015), and the reaction was carried out following manufacturer’s protocol. cDNA was synthesized using SuperScript III Reverse Transcriptase as described above.

### qPCR

Multiplex qPCR was used to measure LTR, *gag* and *env* expression. The reactions were performed using TaqMan™ Fast Advanced Master Mix (Thermo Fischer Scientific cat. 4444555). Primers and probes for the HIV-1 transcripts LTR, *gag* and *env* are described in Supplementary table 5. Primers and probes were used at a final concentration of 0.25 µM. A standard curve for quantification was based on 10-fold serial dilutions of a plasmid from HIV-1 molecular clone HIV-1 NL4-3 NIH (HIV Reagent Program cat. ARP-114) calculated by Design and Analysis software (version 2.6.0 Thermo Fischer Scientific). For relative expression analysis, host gene *PPiA* was used for normalization, quantified using a predesigned PrimeTime qPCR Assay (Hs.PT.58v.38887593.g, Integrated DNA Technologies). Expression was calculated using the formula 2^-(*Ct*_*target*_-*Ct*_*ref*_)^. The frequency of cells expressing HIV-1 transcripts was determined using Most Probable Number (MPN) analysis based on the proportion of wells testing positive at each cell dilution in the limiting dilution assay. Statistical analysis employed a custom qPCR Poisson Distribution Analysis Shiny application (qpcrpoissondist), which uses the MPN R package (v0.4.0) to perform maximum likelihood estimation of microbial densities from serial dilution data. This method calculates the most probable number of target templates per sample assuming a Poisson distribution, maximizing the likelihood function to estimate λ (lambda), the mean number of template copies per reaction. The distribution of positive and negative wells across dilution series informs this estimation, where the probability of detection is given by P(detection) = 1 − e^−λ^, representing the chance of detecting at least one template copy per well. For wells with detectable HIV-1 RNA, copy numbers per expressing cell were calculated by dividing the total RNA copies detected by the estimated number of expressing cells in that well, as established from the MPN-based frequency analysis. This approach accounts for the stochastic distribution of expressing cells across dilution series and provides quantitative estimates with confidence intervals both for the proportion of cells capable of HIV-1 expression and for the transcriptional output per expressing cell under different treatment conditions.

### Infection of healthy donor cells

The presence of infectious particles in the supernatant of the cultured cells was evaluated by incubation with healthy donor CD4+ T cells. CD4+ T cells were isolated from PBMCs from healthy donors using magnetic separation (Miltenyi cat. 130-096-533) and cultured in 96 well U bottom plates, at 5 x 10^5^ cells/well, in activation media. After 24 hours, the culture supernatants were exchanged for supernatants of resting or activated HIV-1 CD4+ T cell clones cultured in the absence of ARVs for 24 hours. Cells were incubated for 6 days at 37°C with 5% CO_2_ and supernatants were tested for Gag p24 by Elisa using Lenti-X™ p24 Rapid Titer Kit (Takara cat. 631476), according to manufacturer’s instructions.

### Single-cell RNA sequencing

Single-cell mRNA sequencing was performed using the 10X genomics platform. Chromium Single Cell Library & Gel Bead Kit (10X Genomics cat. PN-1000014) and Chromium Single Cell V(D)J Enrichment Kit, Human T Cell (10X Genomics cat. PN-1000005) were used to create the gene expression and V(D)J libraries, respectively. Libraries were prepared following the 10X genomics protocol and sequenced in the NovaSeq 6000 System (Illumina) using a SP Flow cell (500 cycles) (Illumina cat. 20028402).

Binary base call (BCL) files were demultiplexed and BCLtoFastq software (Illumina) was used to transform the files to FASTQ format. Cellranger multi (v8.0.1 Illumina) was used to align the reads with a modified version of human genome hg38 (Perez et al., 2024), that includes HIV specific sequences from each HIV-1 provirus clone. Seurat (v5.3.0) was used for the analysis using Rstudio server (2024.12.0 Build 467). Cells outside the 200 to 2,500 range and/or with mitochondrial content higher then 10% were filtered out. SCTransform was used to merge sample batches and also to normalize and scale. Cellranger multi was also used to assemble and annotate TCR sequences using 10x VDJ human reference (GRCh38-alts-ensembl-7.1.0). Filtering and analysis of the resulting contig annotations was done using R studio. Latent clones were determined based on identical combined TRA CDR3 and TRB CDR3 nucleotide sequences, utilizing TCR sequence information obtained either previously (603, 5104, 5125, 9247 and B207)^4^ or during this study (5203 and P301).

### Mapping scRNA-seq data from cultured cells to a CD4**⁺** T cell reference

The CD4+ T cell population was extracted from a publicly available, multimodally annotated human peripheral blood reference dataset (Hao et al., 2021). A UMAP reference was reconstructed using the first 50 principal components derived from the RNA expression slot. Cultured cells from each individual were then anchored and mapped onto this reference using Seurat’s FindTransferAnchors and MapQuery functions (reference.reduction = "pca", dims = 1:50, reduction.model = umap).

### Mapping scRNA-seq data from cultured cells onto the published UMAP reference

Single-cell RNA-seq data from cultured cells were mapped onto a previously published UMAP of the same participants (Weymar et al., 2022), which served as the reference. Mapping was performed using the FindTransferAnchors and MapQuery functions from Seurat, with parameters reference.reduction = "pca", dims = 1:30, and reduction.model = umap. Cell type labels were transferred using the refdata = clusters argument. Only cells with a label prediction score ≥ 0.95 were retained for downstream analysis.

### Differential expression analysis

We tested for differential expression using the Seurat function FindMarkers() with default parameters, except min.pct = 0. Genes with an absolute average log2 fold change ≥ 1 and adjusted p-value < 0.05 were considered differentially expressed (DEGs). For comparisons between HIV-positive and HIV-negative cells, latent cells were stratified based on 5’LTR expression. Cells with SCT-normalized expression > 0.01 were labeled as HIV-positive, while those below this threshold were classified as HIV-negative. Upregulated overlapping DEGs across different clones were selected if detected in at least two clones under resting or activated conditions. For comparisons between HIV-infected and non-infected cells, clone-specific HIV-infected cells were compared to non-infected cells pooled across all clones. In this case, upregulated overlapping DEGs were selected if detected in a minimum of 10% of cells from at least four clones under resting or activated conditions.

## Acknowledgments

We thank all study participants who devoted time to our research, the Rockefeller University Hospital Research support office and nursing staff, all members of the Nussenzweig laboratory for discussions, and M. Jankovic and Tacio Waldetario for laboratory support. We thank C. Zhao, H. Duan, C. Lai, and S. Huang from the Genomics Resource Center at the Rockefeller University for preparing and sequencing the 10x Genomics. We also thank J.P. Truman and K.M. Gordon for operating the cell sorters. This work was supported by the National Institutes of Health (grants UM1 AI100663 and R01AI129795 to M.C.Nussenzweig), (grants UM1AI191237 and UM1AI164565, the latter also supported by NIMH, NIDA, NINDS, NIDDK, NHLBI to R.B.Jones), REACH Delaney (grant UM1 AI164565 to M.Caskey), the Einstein-Rockefeller-CUNY Center for AIDS Research (grant 1P30AI124414-01A1), BEAT-HIV Delaney (grant UM1 AI126620 to M. Caskey), the Bill and Melinda Gates Foundation (INV-008540 and INV-002705) and the Stavros Niarchos Foundation (SNF) as part of its grant to the SNF Institute for Global Infectious Disease Research at The Rockefeller University. K.Lenart is supported by Swedish Research Council grant 2024-00448.

M.C.Nussenzweig is a Howard Hughes Medical Institute (HHMI) Investigator. This article is subject to HHMI’s Open Access to Publications policy. HHMI lab heads have previously granted a non-exclusive CC BY 4.0 license to the public and a sublicensable license to HHMI in their research articles. Pursuant to those licenses, the author-accepted manuscript of this article can be made freely available under a CC BY 4.0 license immediately upon publication.

M.C. Nussenzweig had a patent on anti-HIV-1 antibodies 3BNC117 and 10-1074 licensed to Gilead and a patent to C144 and C135 licensed to Bristol Meyers Squib. The authors have no conflicting financial interests.

## Figure legends

**Supplementary figure 1.**
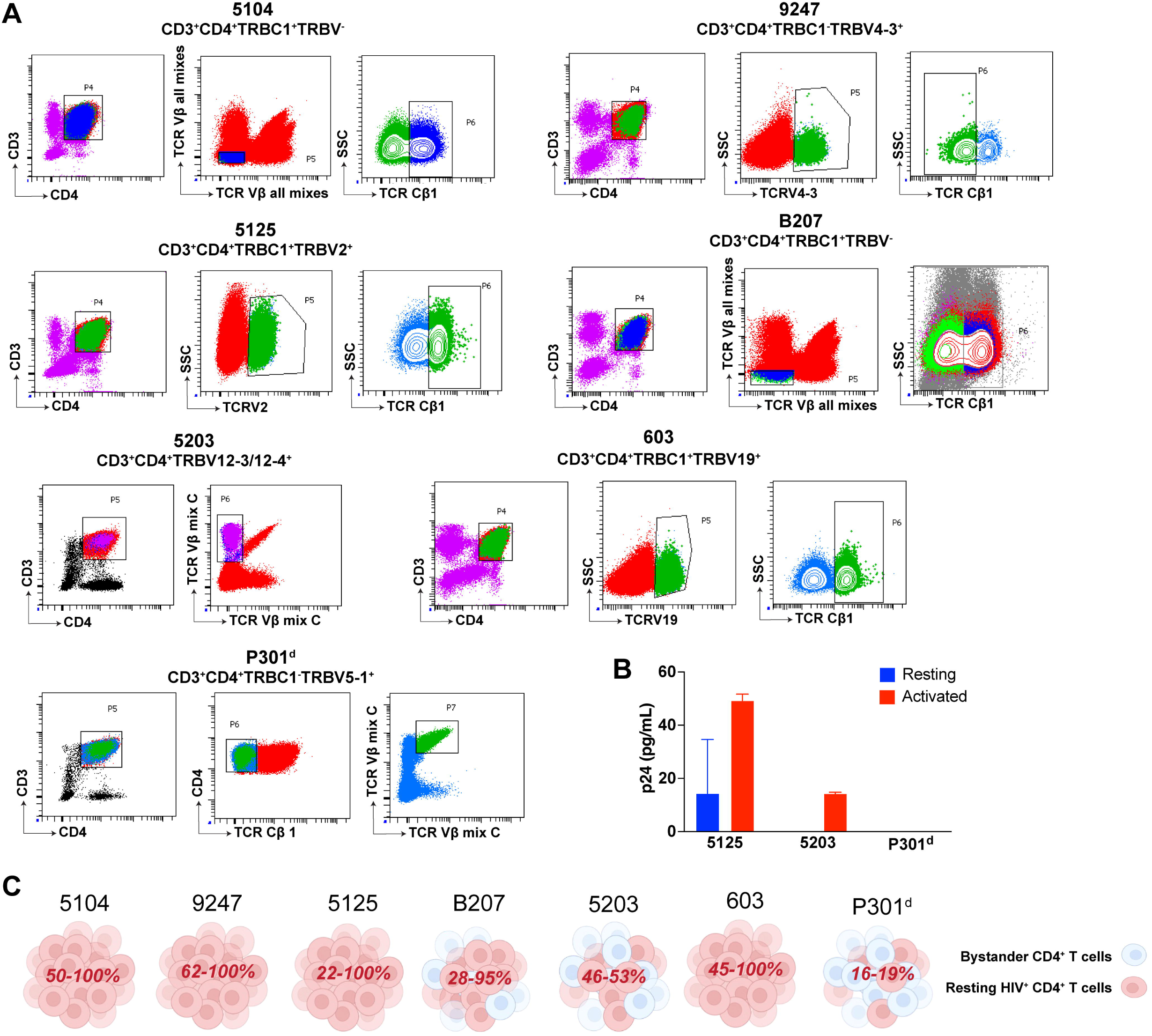
Enrichment of HIV-1+ cells. **(A)** Enrichment of each HIV-1 clone by cell sorting. (**B)** p24 detection by Elisa in the supernatant of healthy cells infected with supernatant from the latent cell clones. **(C)** Diagram showing final enrichment for each specific clone.

**Supplementary figure 2.**
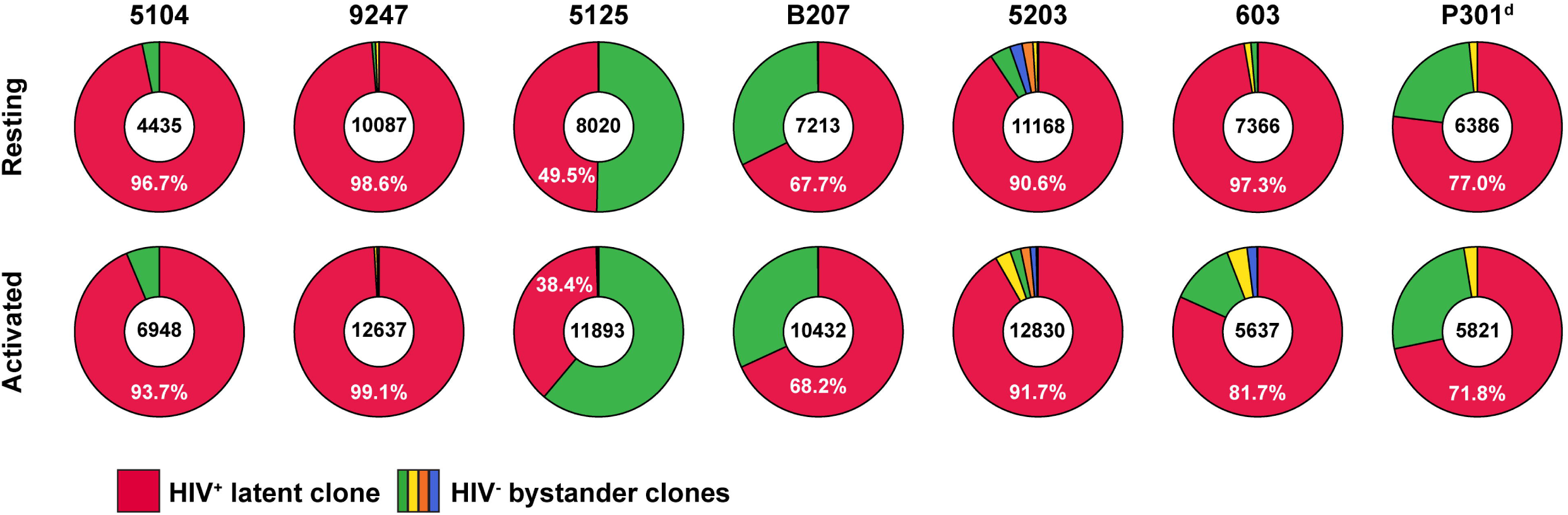
Number of cells analyzed by single-cell RNA sequencing (10X). Pie charts represent the clonal distribution of cells used for single-cell RNA analysis in resting and activated conditions. Red slices represent the HIV-1+ CD4 T cell clones, with frequency shown in white. HIV-1 negative clones are presented in green, yellow, orange and blue.

**Supplementary figure 3.**
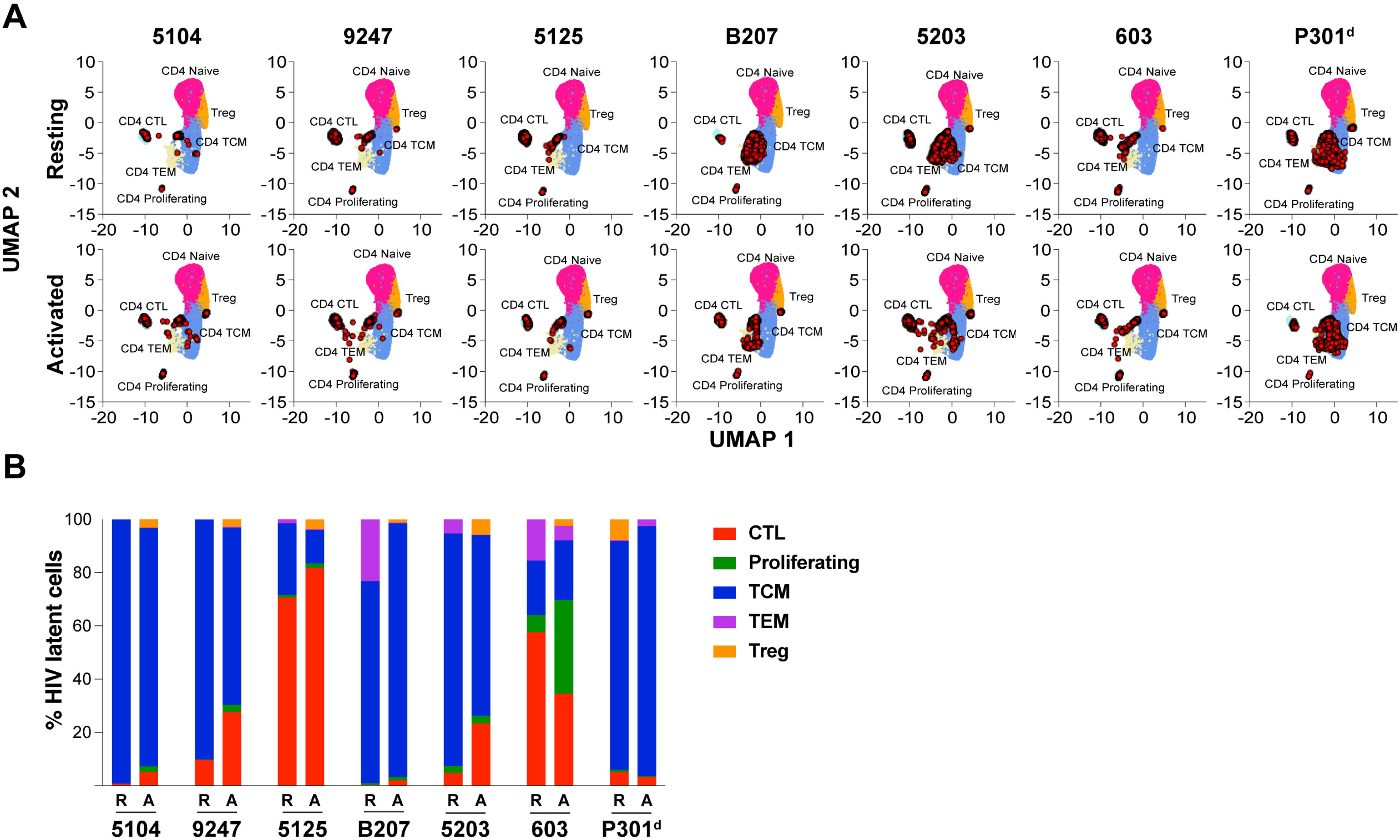
Projection of single-cell gene expression data on a HIV-1 CD4+ T cell UMAP. **(A)** Projection of the RNA expression data of 41,001cells from the HIV-1+ CD4 T cell clones on a multimodal UMAP of CD4+ T cells from HIV-negative donors, in resting and activated states. HIV-1+ CD4 T cell clones are represented as red dots. **(B)** Distribution of the CD4+ T cells subpopulation, based on the UMAP projection, for each clone, in resting (R) and activated (A) conditions.

**Supplementary figure 4.**
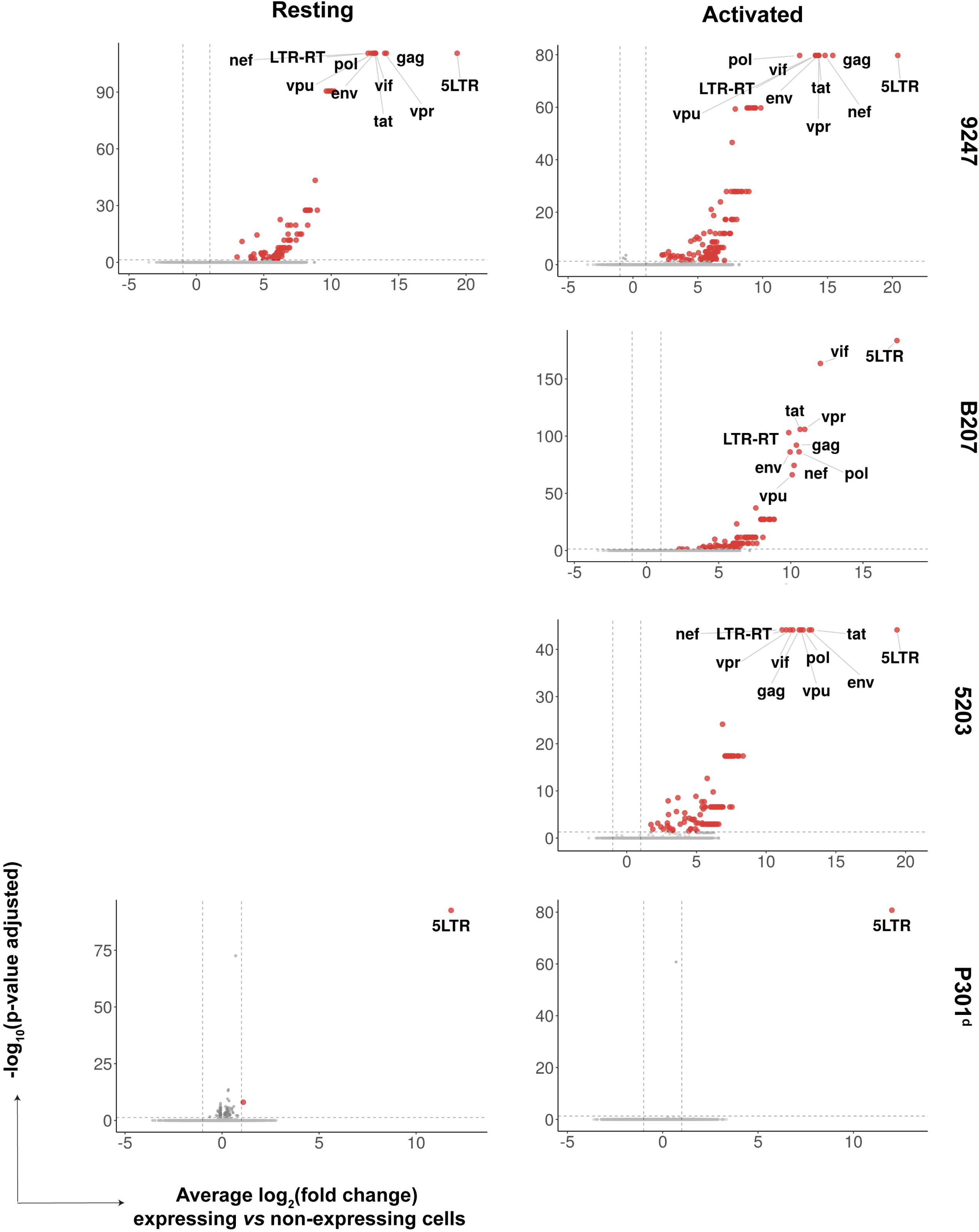
Differential expression between cells expressing HIV-1 transcripts and latent HIV-positive cells. Volcano plots showing differentially expressed genes between HIV-expressing and non-expressing cells from clones 9247, B207,5203 and P301^d^ under resting or activated conditions. For clones B207 and 5203 in resting and clone 603 the number of expressing cells was not sufficient for this analysis.

## Supplementary tables

**Supplementary table 1. Integration site.** Information of the integration site for each clone. Detailed information on the position of the integration within the host genome is provided. Integration site sequence is also provided.

**Supplementary table 2. Total number of cells analyzed by single-cell RNA sequencing.** The number and frequency of latent (HIV+) and non-latent (HIV-) cells are presented. Data for resting and activated conditions are presented in separate tabs.

**Supplementary table 3. Differential expression between HIV-positive and HIV-negative CD4+ T cell.** Each tab includes the list of genes that are differentially expressed, for each clone in resting and activated conditions.

**Supplementary table 4. Proportion of cells expressing HIV-1 LTR based on single-cell RNA sequencing.** Percentage of cells expressing HIV-1 LTR transcripts for each clone, in resting and activated states, according to single-cell RNA sequencing (10X) analysis.

**Supplementary table 5. List of primers and probes.** Primers and probes used throughout this work.

## References

Archin, N.M., A.L. Liberty, A.D. Kashuba, S.K. Choudhary, J.D. Kuruc, A.M. Crooks, D.C. Parker, E.M. Anderson, M.F. Kearney, M.C. Strain, D.D. Richman, M.G. Hudgens, R.J. Bosch, J.M. Coffin, J.J. Eron, D.J. Hazuda, and D.M. Margolis. 2012. Administration of vorinostat disrupts HIV-1 latency in patients on antiretroviral therapy. Nature 487:482–485.

Armani-Tourret, M., C. Gao, C.A. Hartana, W. Sun, L. Carrere, L. Vela, A. Hochroth, M. Bellefroid, A. Sbrolla, K. Shea, T. Flynn, I. Roseto, Y. Rassadkina, C. Lee, F. Giguel, R. Malhotra, F.D. Bushman, R.T. Gandhi, X.G. Yu, D.R. Kuritzkes, and M. Lichterfeld. 2024. Selection of epigenetically privileged HIV-1 proviruses during treatment with panobinostat and interferon-alpha2a. Cell 187:1238–1254 e1214.

Bachmann, N., C. von Siebenthal, V. Vongrad, T. Turk, K. Neumann, N. Beerenwinkel, J. Bogojeska, J. Fellay, V. Roth, Y.L. Kok, C.W. Thorball, A. Borghesi, S. Parbhoo, M. Wieser, J. Boni, M. Perreau, T. Klimkait, S. Yerly, M. Battegay, A. Rauch, M. Hoffmann, E. Bernasconi, M. Cavassini, R.D. Kouyos, H.F. Gunthard, K.J. Metzner, and H.I.V.C.S. Swiss. 2019. Determinants of HIV-1 reservoir size and long-term dynamics during suppressive ART. Nat Commun 10:3193.

Battivelli, E., M.S. Dahabieh, M. Abdel-Mohsen, J.P. Svensson, I. Tojal Da Silva, L.B. Cohn, A. Gramatica, S. Deeks, W.C. Greene, S.K. Pillai, and E. Verdin. 2018. Distinct chromatin functional states correlate with HIV latency reactivation in infected primary CD4(+) T cells. Elife 7:

Bertagnolli, L.N., J. Varriale, S. Sweet, J. Brockhurst, F.R. Simonetti, J. White, S. Beg, K. Lynn, K. Mounzer, I. Frank, P. Tebas, K.J. Bar, L.J. Montaner, R.F. Siliciano, and J.D. Siliciano. 2020. Autologous IgG antibodies block outgrowth of a substantial but variable fraction of viruses in the latent reservoir for HIV-1. Proc Natl Acad Sci U S A 117:32066–32077.

Brouwer, I., and T.L. Lenstra. 2019. Visualizing transcription: key to understanding gene expression dynamics. Curr Opin Chem Biol 51:122–129.

Bruner, K.M., Z. Wang, F.R. Simonetti, A.M. Bender, K.J. Kwon, S. Sengupta, E.J. Fray, S.A. Beg, A.A.R. Antar, K.M. Jenike, L.N. Bertagnolli, A.A. Capoferri, J.T. Kufera, A. Timmons, C. Nobles, J. Gregg, N. Wada, Y.C. Ho, H. Zhang, J.B. Margolick, J.N. Blankson, S.G. Deeks, F.D. Bushman, J.D. Siliciano, G.M. Laird, and R.F. Siliciano. 2019. A quantitative approach for measuring the reservoir of latent HIV-1 proviruses. Nature 566:120–125.

Bui, J.K., E.K. Halvas, E. Fyne, M.D. Sobolewski, D. Koontz, W. Shao, B. Luke, F.F. Hong, M.F. Kearney, and J.W. Mellors. 2017a. Ex vivo activation of CD4+ T-cells from donors on suppressive ART can lead to sustained production of infectious HIV-1 from a subset of infected cells. PLoS Pathog 13:e1006230.

Bui, J.K., M.D. Sobolewski, B.F. Keele, J. Spindler, A. Musick, A. Wiegand, B.T. Luke, W. Shao, S.H. Hughes, J.M. Coffin, M.F. Kearney, and J.W. Mellors. 2017b. Proviruses with identical sequences comprise a large fraction of the replication-competent HIV reservoir. PLoS Pathog 13:e1006283.

Cano-Gamez, E., B. Soskic, T.I. Roumeliotis, E. So, D.J. Smyth, M. Baldrighi, D. Wille, N. Nakic, J. Esparza-Gordillo, C.G.C. Larminie, P.G. Bronson, D.F. Tough, W.C. Rowan, J.S. Choudhary, and G. Trynka. 2020. Single-cell transcriptomics identifies an effectorness gradient shaping the response of CD4(+) T cells to cytokines. Nat Commun 11:1801.

Cho, A., C. Gaebler, T. Olveira, V. Ramos, M. Saad, J.C.C. Lorenzi, A. Gazumyan, S. Moir, M. Caskey, T.W. Chun, and M.C. Nussenzweig. 2022. Longitudinal clonal dynamics of HIV-1 latent reservoirs measured by combination quadruplex polymerase chain reaction and sequencing. Proc Natl Acad Sci U S A 119:

Chun, T.W., D. Engel, M.M. Berrey, T. Shea, L. Corey, and A.S. Fauci. 1998a. Early establishment of a pool of latently infected, resting CD4(+) T cells during primary HIV-1 infection. Proc Natl Acad Sci U S A 95:8869–8873.

Chun, T.W., D. Engel, S.B. Mizell, L.A. Ehler, and A.S. Fauci. 1998b. Induction of HIV-1 replication in latently infected CD4+ T cells using a combination of cytokines. J Exp Med 188:83–91.

Cohn, L.B., N. Chomont, and S.G. Deeks. 2020. The Biology of the HIV-1 Latent Reservoir and Implications for Cure Strategies. Cell Host Microbe 27:519–530.

Cohn, L.B., I.T. da Silva, R. Valieris, A.S. Huang, J.C.C. Lorenzi, Y.Z. Cohen, J.A. Pai, A.L. Butler, M. Caskey, M. Jankovic, and M.C. Nussenzweig. 2018. Clonal CD4(+) T cells in the HIV-1 latent reservoir display a distinct gene profile upon reactivation. Nat Med 24:604–609.

Cohn, L.B., I.T. Silva, T.Y. Oliveira, R.A. Rosales, E.H. Parrish, G.H. Learn, B.H. Hahn, J.L. Czartoski, M.J. McElrath, C. Lehmann, F. Klein, M. Caskey, B.D. Walker, J.D. Siliciano, R.F. Siliciano, M. Jankovic, and M.C. Nussenzweig. 2015. HIV-1 integration landscape during latent and active infection. Cell 160:420–432.

Collora, J.A., and Y.C. Ho. 2023. Integration site-dependent HIV-1 promoter activity shapes host chromatin conformation. Genome Res 33:891–906.

Collora, J.A., R. Liu, D. Pinto-Santini, N. Ravindra, C. Ganoza, J.R. Lama, R. Alfaro, J. Chiarella, S. Spudich, K. Mounzer, P. Tebas, L.J. Montaner, D. van Dijk, A. Duerr, and Y.C. Ho. 2022. Single-cell multiomics reveals persistence of HIV-1 in expanded cytotoxic T cell clones. Immunity 55:1013–1031 e1017.

Crooks, A.M., R. Bateson, A.B. Cope, N.P. Dahl, M.K. Griggs, J.D. Kuruc, C.L. Gay, J.J. Eron, D.M. Margolis, R.J. Bosch, and N.M. Archin. 2015. Precise Quantitation of the Latent HIV-1 Reservoir: Implications for Eradication Strategies. J Infect Dis 212:1361–1365.

Damour, A., V. Slaninova, O. Radulescu, E. Bertrand, and E. Basyuk. 2023. Transcriptional Stochasticity as a Key Aspect of HIV-1 Latency. Viruses 15:

Davey, R.T., Jr., N. Bhat, C. Yoder, T.W. Chun, J.A. Metcalf, R. Dewar, V. Natarajan, R.A. Lempicki, J.W. Adelsberger, K.D. Miller, J.A. Kovacs, M.A. Polis, R.E. Walker, J. Falloon, H. Masur, D. Gee, M. Baseler, D.S. Dimitrov, A.S. Fauci, and H.C. Lane. 1999. HIV-1 and T cell dynamics after interruption of highly active antiretroviral therapy (HAART) in patients with a history of sustained viral suppression. Proc Natl Acad Sci U S A 96:15109–15114.

Debrabander, Q., K.S. Hensley, C.K. Psomas, W. Bramer, T. Mahmoudi, B.J. van Welzen, A. Verbon, and C. Rokx. 2023. The efficacy and tolerability of latency-reversing agents in reactivating the HIV-1 reservoir in clinical studies: a systematic review. J Virus Erad 9:100342.

Demoustier, A., B. Gubler, O. Lambotte, M.G. de Goer, C. Wallon, C. Goujard, J.F. Delfraissy, and Y. Taoufik. 2002. In patients on prolonged HAART, a significant pool of HIV infected CD4 T cells are HIV-specific. AIDS 16:1749–1754.

Douek, D.C., J.M. Brenchley, M.R. Betts, D.R. Ambrozak, B.J. Hill, Y. Okamoto, J.P. Casazza, J. Kuruppu, K. Kunstman, S. Wolinsky, Z. Grossman, M. Dybul, A. Oxenius, D.A. Price, M. Connors, and R.A. Koup. 2002. HIV preferentially infects HIV-specific CD4+ T cells. Nature 417:95–98.

Dube, M., O. Tastet, C. Dufour, G. Sannier, N. Brassard, G.G. Delgado, A. Pagliuzza, C. Richard, M. Nayrac, J.P. Routy, A. Prat, J.D. Estes, R. Fromentin, N. Chomont, and D.E. Kaufmann. 2023. Spontaneous HIV expression during suppressive ART is associated with the magnitude and function of HIV-specific CD4(+) and CD8(+) T cells. Cell Host Microbe 31:1507–1522 e1505.

Einkauf, K.B., G.Q. Lee, C. Gao, R. Sharaf, X. Sun, S. Hua, S.M. Chen, C. Jiang, X. Lian, F.Z. Chowdhury, E.S. Rosenberg, T.W. Chun, J.Z. Li, X.G. Yu, and M. Lichterfeld. 2019. Intact HIV-1 proviruses accumulate at distinct chromosomal positions during prolonged antiretroviral therapy. J Clin Invest 129:988–998.

Einkauf, K.B., M.R. Osborn, C. Gao, W. Sun, X. Sun, X. Lian, E.M. Parsons, G.T. Gladkov, K.W. Seiger, J.E. Blackmer, C. Jiang, S.A. Yukl, E.S. Rosenberg, X.G. Yu, and M. Lichterfeld. 2022. Parallel analysis of transcription, integration, and sequence of single HIV-1 proviruses. Cell 185:266–282 e215.

Esmaeilzadeh, E., B. Etemad, C.L. Lavine, L. Garneau, Y. Li, J. Regan, C. Wong, R. Sharaf, E. Connick, P. Volberding, M. Sagar, M.S. Seaman, and J.Z. Li. 2023. Autologous neutralizing antibodies increase with early antiretroviral therapy and shape HIV rebound after treatment interruption. Sci Transl Med 15:eabq4490.

Finzi, D., M. Hermankova, T. Pierson, L.M. Carruth, C. Buck, R.E. Chaisson, T.C. Quinn, K. Chadwick, J. Margolick, R. Brookmeyer, J. Gallant, M. Markowitz, D.D. Ho, D.D. Richman, and R.F. Siliciano. 1997. Identification of a reservoir for HIV-1 in patients on highly active antiretroviral therapy. Science 278:1295–1300.

Gaebler, C., L. Nogueira, E. Stoffel, T.Y. Oliveira, G. Breton, K.G. Millard, M. Turroja, A. Butler, V. Ramos, M.S. Seaman, J.D. Reeves, C.J. Petroupoulos, I. Shimeliovich, A. Gazumyan, C.S. Jiang, N. Jilg, J.F. Scheid, R. Gandhi, B.D. Walker, M.C. Sneller, A. Fauci, T.W. Chun, M. Caskey, and M.C. Nussenzweig. 2022. Prolonged viral suppression with anti-HIV-1 antibody therapy. Nature 606:368–374.

Gunst, J.D., J.F. Hojen, and O.S. Sogaard. 2020. Broadly neutralizing antibodies combined with latency-reversing agents or immune modulators as strategy for HIV-1 remission. Curr Opin HIV AIDS 15:309–315.

Hao, Y., S. Hao, E. Andersen-Nissen, W.M. Mauck, S. Zheng, A. Butler, M.J. Lee, A.J. Wilk, C. Darby, M. Zager, P. Hoffman, M. Stoeckius, E. Papalexi, E.P. Mimitou, J. Jain, A. Srivastava, T. Stuart, L.M. Fleming, B. Yeung, A.J. Rogers, J.M. McElrath, C.A. Blish, R. Gottardo, P. Smibert, and R. Satija. 2021. Integrated analysis of multimodal single-cell data. Cell 184:3573–3587.e3529.

Ho, Y.C., L. Shan, N.N. Hosmane, J. Wang, S.B. Laskey, D.I. Rosenbloom, J. Lai, J.N. Blankson, J.D. Siliciano, and R.F. Siliciano. 2013. Replication-competent noninduced proviruses in the latent reservoir increase barrier to HIV-1 cure. Cell 155:540–551.

Horsburgh, B.A., E. Lee, B. Hiener, J.S. Eden, T.E. Schlub, S. von Stockenstrom, L. Odevall, J.M. Milush, T. Liegler, E. Sinclair, R. Hoh, E.A. Boritz, D.C. Douek, R. Fromentin, N. Chomont, S.G. Deeks, F.M. Hecht, and S. Palmer. 2020. High levels of genetically intact HIV in HLA-DR+ memory T cells indicates their value for reservoir studies. AIDS 34:659–668.

Hosmane, N.N., K.J. Kwon, K.M. Bruner, A.A. Capoferri, S. Beg, D.I. Rosenbloom, B.F. Keele, Y.C. Ho, J.D. Siliciano, and R.F. Siliciano. 2017. Proliferation of latently infected CD4(+) T cells carrying replication-competent HIV-1: Potential role in latent reservoir dynamics. J Exp Med 214:959–972.

Huang, A.S., V. Ramos, T.Y. Oliveira, C. Gaebler, M. Jankovic, M.C. Nussenzweig, and L.B. Cohn. 2021. Integration features of intact latent HIV-1 in CD4+ T cell clones contribute to viral persistence. J Exp Med 218:

Jiang, C., X. Lian, C. Gao, X. Sun, K.B. Einkauf, J.M. Chevalier, S.M.Y. Chen, S. Hua, B. Rhee, K. Chang, J.E. Blackmer, M. Osborn, M.J. Peluso, R. Hoh, M. Somsouk, J. Milush, L.N. Bertagnolli, S.E. Sweet, J.A. Varriale, P.D. Burbelo, T.W. Chun, G.M. Laird, E. Serrao, A.N. Engelman, M. Carrington, R.F. Siliciano, J.M. Siliciano, S.G. Deeks, B.D. Walker, M. Lichterfeld, and X.G. Yu. 2020. Distinct viral reservoirs in individuals with spontaneous control of HIV-1. Nature 585:261–267.

Jordan, A., D. Bisgrove, and E. Verdin. 2003. HIV reproducibly establishes a latent infection after acute infection of T cells in vitro. EMBO J 22:1868–1877.

Jordan, A., P. Defechereux, and E. Verdin. 2001. The site of HIV-1 integration in the human genome determines basal transcriptional activity and response to Tat transactivation. EMBO J 20:1726–1738.

Lewinski, M.K., D. Bisgrove, P. Shinn, H. Chen, C. Hoffmann, S. Hannenhalli, E. Verdin, C.C. Berry, J.R. Ecker, and F.D. Bushman. 2005. Genome-wide analysis of chromosomal features repressing human immunodeficiency virus transcription. J Virol 79:6610–6619.

Lian, X., K.W. Seiger, E.M. Parsons, C. Gao, W. Sun, G.T. Gladkov, I.C. Roseto, K.B. Einkauf, M.R. Osborn, J.M. Chevalier, C. Jiang, J. Blackmer, M. Carrington, E.S. Rosenberg, M.M. Lederman, D.K. McMahon, R.J. Bosch, J.M. Jacobson, R.T. Gandhi, M.J. Peluso, T.W. Chun, S.G. Deeks, X.G. Yu, and M. Lichterfeld. 2023. Progressive transformation of the HIV-1 reservoir cell profile over two decades of antiviral therapy. Cell Host Microbe 31:83–96 e85.

Lorenzi, J.C., Y.Z. Cohen, L.B. Cohn, E.F. Kreider, J.P. Barton, G.H. Learn, T. Oliveira, C.L. Lavine, J.A. Horwitz, A. Settler, M. Jankovic, M.S. Seaman, A.K. Chakraborty, B.H. Hahn, M. Caskey, and M.C. Nussenzweig. 2016. Paired quantitative and qualitative assessment of the replication-competent HIV-1 reservoir and comparison with integrated proviral DNA. Proc Natl Acad Sci U S A 113:E7908–E7916.

Lu, C.L., J.A. Pai, L. Nogueira, P. Mendoza, H. Gruell, T.Y. Oliveira, J. Barton, J.C.C. Lorenzi, Y.Z. Cohen, L.B. Cohn, F. Klein, M. Caskey, M.C. Nussenzweig, and M. Jankovic. 2018. Relationship between intact HIV-1 proviruses in circulating CD4(+) T cells and rebound viruses emerging during treatment interruption. Proc Natl Acad Sci U S A 115:E11341–E11348.

Margolis, D.M., and N.M. Archin. 2017. Proviral Latency, Persistent Human Immunodeficiency Virus Infection, and the Development of Latency Reversing Agents. J Infect Dis 215:S111–S118.

McMyn, N.F., J. Varriale, E.J. Fray, C. Zitzmann, H. MacLeod, J. Lai, A. Singhal, M. Moskovljevic, M.A. Garcia, B.M. Lopez, V. Hariharan, K. Rhodehouse, K. Lynn, P. Tebas, K. Mounzer, L.J. Montaner, E. Benko, C. Kovacs, R. Hoh, F.R. Simonetti, G.M. Laird, S.G. Deeks, R.M. Ribeiro, A.S. Perelson, R.F. Siliciano, and J.M. Siliciano. 2023. The latent reservoir of inducible, infectious HIV-1 does not decrease despite decades of antiretroviral therapy. J Clin Invest 133:

Mendoza, P., H. Gruell, L. Nogueira, J.A. Pai, A.L. Butler, K. Millard, C. Lehmann, I. Suarez, T.Y. Oliveira, J.C.C. Lorenzi, Y.Z. Cohen, C. Wyen, T. Kummerle, T. Karagounis, C.L. Lu, L. Handl, C. Unson-O’Brien, R. Patel, C. Ruping, M. Schlotz, M. Witmer-Pack, I. Shimeliovich, G. Kremer, E. Thomas, K.E. Seaton, J. Horowitz, A.P. West, Jr., P.J. Bjorkman, G.D. Tomaras, R.M. Gulick, N. Pfeifer, G. Fatkenheuer, M.S. Seaman, F. Klein, M. Caskey, and M.C. Nussenzweig. 2018. Combination therapy with anti-HIV-1 antibodies maintains viral suppression. Nature 561:479–484.

Niessl, J., A.E. Baxter, P. Mendoza, M. Jankovic, Y.Z. Cohen, A.L. Butler, C.L. Lu, M. Dube, I. Shimeliovich, H. Gruell, F. Klein, M. Caskey, M.C. Nussenzweig, and D.E. Kaufmann. 2020. Combination anti-HIV-1 antibody therapy is associated with increased virus-specific T cell immunity. Nat Med 26:222–227.

Olesen, R., S. Vigano, T.A. Rasmussen, O.S. Sogaard, Z. Ouyang, M. Buzon, A. Bashirova, M. Carrington, S. Palmer, C.R. Brinkmann, X.G. Yu, L. Ostergaard, M. Tolstrup, and M. Lichterfeld. 2015. Innate Immune Activity Correlates with CD4 T Cell-Associated HIV-1 DNA Decline during Latency-Reversing Treatment with Panobinostat. J Virol 89:10176–10189.

Peluso, M.J., P. Bacchetti, K.D. Ritter, S. Beg, J. Lai, J.N. Martin, P.W. Hunt, T.J. Henrich, J.D. Siliciano, R.F. Siliciano, G.M. Laird, and S.G. Deeks. 2020. Differential decay of intact and defective proviral DNA in HIV-1-infected individuals on suppressive antiretroviral therapy. JCI Insight 5:

Perea-Resa, C., and M.D. Blower. 2018. Centromere Biology: Transcription Goes on Stage. Mol Cell Biol 38:

Perez, G., Galt P. Barber, A. Benet-Pages, J. Casper, H. Clawson, M. Diekhans, C. Fischer, Jairo N. Gonzalez, Angie S. Hinrichs, Christopher M. Lee, Luis R. Nassar, Brian J. Raney, Matthew L. Speir, Marijke J. van Baren, Charles J. Vaske, D. Haussler, W.J. Kent, and M. Haeussler. 2024. The UCSC Genome Browser database: 2025 update. Nucleic Acids Research 53:D1243–D1249.

Procopio, F.A., R. Fromentin, D.A. Kulpa, J.H. Brehm, A.G. Bebin, M.C. Strain, D.D. Richman, U. O’Doherty, S. Palmer, F.M. Hecht, R. Hoh, R.J. Barnard, M.D. Miller, D.J. Hazuda, S.G. Deeks, R.P. Sekaly, and N. Chomont. 2015. A Novel Assay to Measure the Magnitude of the Inducible Viral Reservoir in HIV-infected Individuals. EBioMedicine 2:874–883.

Rasmussen, T.A., M. Tolstrup, C.R. Brinkmann, R. Olesen, C. Erikstrup, A. Solomon, A. Winckelmann, S. Palmer, C. Dinarello, M. Buzon, M. Lichterfeld, S.R. Lewin, L. Ostergaard, and O.S. Sogaard. 2014. Panobinostat, a histone deacetylase inhibitor, for latent-virus reactivation in HIV-infected patients on suppressive antiretroviral therapy: a phase 1/2, single group, clinical trial. Lancet HIV 1:e13–21.

Reeves, D.B., C. Gaebler, T.Y. Oliveira, M.J. Peluso, J.T. Schiffer, L.B. Cohn, S.G. Deeks, and M.C. Nussenzweig. 2023. Impact of misclassified defective proviruses on HIV reservoir measurements. Nat Commun 14:4186.

Rodari, A., G. Darcis, and C.M. Van Lint. 2021. The Current Status of Latency Reversing Agents for HIV-1 Remission. Annu Rev Virol 8:491–514.

Ruelas, D.S., and W.C. Greene. 2013. An integrated overview of HIV-1 latency. Cell 155:519–529.

Siliciano, J.D., J. Kajdas, D. Finzi, T.C. Quinn, K. Chadwick, J.B. Margolick, C. Kovacs, S.J. Gange, and R.F. Siliciano. 2003. Long-term follow-up studies confirm the stability of the latent reservoir for HIV-1 in resting CD4+ T cells. Nat Med 9:727–728.

Siliciano, J.D., and R.F. Siliciano. 2022. In Vivo Dynamics of the Latent Reservoir for HIV-1: New Insights and Implications for Cure. Annu Rev Pathol 17:271–294.

Simonetti, F.R., M.D. Sobolewski, E. Fyne, W. Shao, J. Spindler, J. Hattori, E.M. Anderson, S.A. Watters, S. Hill, X. Wu, D. Wells, L. Su, B.T. Luke, E.K. Halvas, G. Besson, K.J. Penrose, Z. Yang, R.W. Kwan, C. Van Waes, T. Uldrick, D.E. Citrin, J. Kovacs, M.A. Polis, C.A. Rehm, R. Gorelick, M. Piatak, B.F. Keele, M.F. Kearney, J.M. Coffin, S.H. Hughes, J.W. Mellors, and F. Maldarelli. 2016. Clonally expanded CD4+ T cells can produce infectious HIV-1 in vivo. Proc Natl Acad Sci U S A 113:1883–1888.

Simonetti, F.R., H. Zhang, G.P. Soroosh, J. Duan, K. Rhodehouse, A.L. Hill, S.A. Beg, K. McCormick, H.E. Raymond, C.L. Nobles, J.K. Everett, K.J. Kwon, J.A. White, J. Lai, J.B. Margolick, R. Hoh, S.G. Deeks, F.D. Bushman, J.D. Siliciano, and R.F. Siliciano. 2021. Antigen-driven clonal selection shapes the persistence of HIV-1-infected CD4+ T cells in vivo. J Clin Invest 131:

Sogaard, O.S., M.E. Graversen, S. Leth, R. Olesen, C.R. Brinkmann, S.K. Nissen, A.S. Kjaer, M.H. Schleimann, P.W. Denton, W.J. Hey-Cunningham, K.K. Koelsch, G. Pantaleo, K. Krogsgaard, M. Sommerfelt, R. Fromentin, N. Chomont, T.A. Rasmussen, L. Ostergaard, and M. Tolstrup. 2015. The Depsipeptide Romidepsin Reverses HIV-1 Latency In Vivo. PLoS Pathog 11:e1005142.

Sun, W., C. Gao, C.A. Hartana, M.R. Osborn, K.B. Einkauf, X. Lian, B. Bone, N. Bonheur, T.W. Chun, E.S. Rosenberg, B.D. Walker, X.G. Yu, and M. Lichterfeld. 2023. Phenotypic signatures of immune selection in HIV-1 reservoir cells. Nature 614:309–317.

Tanaka, K., Y. Kim, M. Roche, and S.R. Lewin. 2022. The role of latency reversal in HIV cure strategies. J Med Primatol 51:278–283.

Tantale, K., E. Garcia-Oliver, M.C. Robert, A. L’Hostis, Y. Yang, N. Tsanov, R. Topno, T. Gostan, A. Kozulic-Pirher, M. Basu-Shrivastava, K. Mukherjee, V. Slaninova, J.C. Andrau, F. Mueller, E. Basyuk, O. Radulescu, and E. Bertrand. 2021. Stochastic pausing at latent HIV-1 promoters generates transcriptional bursting. Nat Commun 12:4503.

Teixeira, A.R., C. Bittar, G.S. Silva Santos, T.Y. Oliveira, A.S. Huang, N. Linden, I. Ferreira, T. Murdza, F. Muecksch, R.B. Jones, M. Caskey, M. Jankovic, and M.C. Nussenzweig. 2024. Transcription of HIV-1 at sites of intact latent provirus integration. J Exp Med 221:

Wagner, T.A., S. McLaughlin, K. Garg, C.Y.K. Cheung, B.B. Larsen, S. Styrchak, H.C. Huang, P.T. Edlefsen, J.I. Mullins, and L.M. Frenkel. 2014. Proliferation of cells with HIV integrated into cancer genes contributes to persistent infection. Science 345:570–573.

Wei, Y., T.C. Davenport, J.A. Collora, H.K. Ma, D. Pinto-Santini, J. Lama, R. Alfaro, A. Duerr, and Y.C. Ho. 2023. Single-cell epigenetic, transcriptional, and protein profiling of latent and active HIV-1 reservoir revealed that IKZF3 promotes HIV-1 persistence. Immunity 56:2584–2601 e2587.

Weinberger, L.S., J.C. Burnett, J.E. Toettcher, A.P. Arkin, and D.V. Schaffer. 2005. Stochastic gene expression in a lentiviral positive-feedback loop: HIV-1 Tat fluctuations drive phenotypic diversity. Cell 122:169–182.

Weymar, G.H.J., Y. Bar-On, T.Y. Oliveira, C. Gaebler, V. Ramos, H. Hartweger, G. Breton, M. Caskey, L.B. Cohn, M. Jankovic, and M.C. Nussenzweig. 2022. Distinct gene expression by expanded clones of quiescent memory CD4(+) T cells harboring intact latent HIV-1 proviruses. Cell Rep 40:111311.

Wiegand, A., J. Spindler, F.F. Hong, W. Shao, J.C. Cyktor, A.R. Cillo, E.K. Halvas, J.M. Coffin, J.W. Mellors, and M.F. Kearney. 2017. Single-cell analysis of HIV-1 transcriptional activity reveals expression of proviruses in expanded clones during ART. Proc Natl Acad Sci U S A 114:E3659–E3668.

Wong, J.K., M. Hezareh, H.F. Gunthard, D.V. Havlir, C.C. Ignacio, C.A. Spina, and D.D. Richman. 1997. Recovery of replication-competent HIV despite prolonged suppression of plasma viremia. Science 278:1291–1295.

Wong, M., Y. Wei, and Y.C. Ho. 2023. Single-cell multiomic understanding of HIV-1 reservoir at epigenetic, transcriptional, and protein levels. Curr Opin HIV AIDS 18:246–256.

Wu, V.H., J.M.L. Nordin, S. Nguyen, J. Joy, F. Mampe, P.M. Del Rio Estrada, F. Torres-Ruiz, M. Gonzalez-Navarro, Y.A. Luna-Villalobos, S. Avila-Rios, G. Reyes-Teran, P. Tebas, L.J. Montaner, K.J. Bar, L.A. Vella, and M.R. Betts. 2023. Profound phenotypic and epigenetic heterogeneity of the HIV-1-infected CD4(+) T cell reservoir. Nat Immunol 24:359–370.

Yukl, S.A., P. Kaiser, P. Kim, S. Telwatte, S.K. Joshi, M. Vu, H. Lampiris, and J.K. Wong. 2018. HIV latency in isolated patient CD4(+) T cells may be due to blocks in HIV transcriptional elongation, completion, and splicing. Sci Transl Med 10:

Zhu, J., Q. Guo, M. Choi, Z. Liang, and K.W.Y. Yuen. 2023. Centromeric and pericentric transcription and transcripts: their intricate relationships, regulation, and functions. Chromosoma 132:211–230.

